# Multi-Parameter Quantitative Imaging of Tumor Microenvironments Reveals Perivascular Immune Niches Associated with Anti-Tumor Immunity

**DOI:** 10.1101/2021.06.17.448881

**Authors:** Caleb R Stoltzfus, Ramya Sivakumar, Leo Kunz, BE Olin Pope, Elena Menietti, Dario Speziale, Roberto Adelfio, Marina Bacac, Sara Colombetti, Mario Perro, Michael Y Gerner

## Abstract

Tumors are populated by a multitude of immune cell types with varied phenotypic and functional properties, which can either promote or inhibit anti-tumor responses. Appropriate localization and function of these cells within tumors is critical for protective immunity, with CD8 T cell infiltration being a biomarker of disease outcome and therapeutic efficacy. Recent multiplexed imaging approaches have revealed highly complex patterns of localization for these immune cell subsets and the generation of distinct tumor microenvironments (TMEs), which can vary among cancer types, individuals, and within individual tumors. While it is recognized that TMEs play a pivotal role in disease progression, a better understanding of their composition, organization, and heterogeneity, as well as how distinct TMEs are reshaped with immunotherapy, is necessary. Here, we performed spatial analysis using multi-parameter confocal imaging, histocytometry, and CytoMAP to study the microanatomical organization of immune cells in two widely used preclinical cancer models, the MC38 colorectal and KPC pancreatic murine tumors engineered to express human carcinoembryonic antigen (CEA). Immune responses were examined in either unperturbed tumors or after immunotherapy with a CEA T cell bispecific (CEA-TCB) surrogate antibody and anti-PD-L1 treatment. CEA-TCB mono and combination immunotherapy markedly enhanced intra-tumoral cellularity of CD8 T cells, dominantly driven by the expansion of TCF1^−^ PD1^+^ effector T cells and with more minor increases in TCF1^+^PD1^+^ resource CD8 T cells. The majority of infiltrating T cells, particularly resource CD8 T cells, were colocalized with dendritic cells (DCs) or activated MHCII^+^ macrophages, but largely avoided the deeper tumor nest regions composed of cancer cells and non-activated macrophages. These myeloid cell – T cell aggregates were found in close proximity to tumor blood vessels, generating perivascular immune niches. This perivascular TME was present in untreated samples and markedly increased after CEA-TCB therapy, with its relative abundance positively associated with response to therapy. Together, these studies demonstrate the utility of advanced spatial analysis in cancer research by revealing that blood vessels are key organizational hubs of innate and adaptive immune cells within tumors, and suggesting the likely relevance of the perivascular immune TME in disease outcome.

## Introduction

Multiplexed imaging and spatially resolved sequencing technologies have revealed complex cellular organization across tissue types and diverse pathological conditions (Gerner et al., 2012; Gerdes et al., 2013; Giesen et al., 2014; Liu et al., 2015; Gerner et al., 2017; Li et al., 2017; Petrovas et al., 2017; Gut et al., 2018; Mao et al., 2018; Lin et al., 2018; Keren et al., 2018; Ak et al., 2019; Vickovic et al., 2019; Eng et al., 2019; Nearchou et al., 2019; Radtke et al., 2020; Schürch et al., 2020; Plumlee et al., 2021; Gern et al., 2021; Leal et al., 2021). Advanced computational and statistical approaches applied to such datasets allow quantification of various spatial properties, such as cellular distance relationships, preferential cell-cell associations, and organization of tissue microenvironments (Caicedo et al., 2017; Schapiro et al., 2017; Coutu et al., 2018; Goltsev et al., 2018; Eling et al., 2020; Schürch et al., 2020; Stoltzfus et al., 2020a; Dries et al., 2021). This in turn allows a data-driven interrogation of how spatial context influences the transcriptional, phenotypic and functional changes within individual cells, and reveals how disorganization of cells can lead to disease pathology. One recent application of such image analytics has been in cancer research, where major efforts are directed at understanding the mechanisms of disease development, as well as the diversity of outcomes in response to immunotherapy.

Tumors are structurally complex tissues, made up of malignant cancer cells and non-malignant host cells, including stromal cells, blood and lymphatic endothelial cells, as well as diverse subsets of immune cells with pro and anti-inflammatory properties, which collectively shape the tumor microenvironment (TME) (Egeblad et al., 2010; Hanahan and Coussens, 2012; Balkwill et al., 2012; Combes et al., 2021). Advanced imaging approaches have demonstrated a high degree of diversity for both cellular composition and spatial organization of cells across different tumor types, among individuals and even within individual tissues, indicating marked heterogeneity and complexity of the TME (Angelo et al., 2014; Giesen et al., 2014; Lee et al., 2017; Wagner et al., 2019; Schürch et al., 2020; Blise et al., 2021). Nevertheless, over a wide variety of samples, specific positional patterns for immune cells have been detected, which correlate with responses to immune therapy. This suggests that microscopy-based technologies may be able to parse out complex cellular patterning in highly heterogeneous tissues and have the potential to serve as powerful tools for companion diagnostics or prognostic studies (Galon et al., 2006; Tsujikawa et al., 2017; Keren et al., 2018; Schürch et al., 2020; Ali et al., 2020; Blise et al., 2021).

Furthermore, imaging has offered invaluable insights into the cellular and molecular mechanisms of disease progression and immune mediated control of tumor growth. Immunologically silent (cold or excluded) tumors lack CD8 T cell infiltration or have T cells excluded to the outer peripheral borders, and frequently exhibit dominant presence of immunosuppressive tumor-associated macrophages and other suppressive myeloid cells within the tumor core. Conversely, potent infiltration of effector CD8 T cells (hot tumors) and numerical dominance over suppressive cells, such as T regulatory cells, has long been appreciated as a hallmark of effective anti-tumor immunity (Tumeh et al., 2014; Beatty et al., 2015; Spranger, 2016; Binnewies et al., 2018; Galon and Bruni, 2019). Differential CD8 T cell infiltration between tumor subtypes is thought to be at least partially regulated by the mutational rates of cancer cells and the subsequent generation of tumor neoantigens (Le et al., 2015; Schumacher and Schreiber, 2015; Yarchoan et al., 2017; Le et al., 2017). As examples, immune infiltrated tumors (e.g., microsatellite instability high colorectal cancer, bladder cancer, melanoma) have been found to be more responsive to checkpoint blockade therapies relative to more immune excluded tumors, such as pancreatic ductal adenocarcinoma or breast cancer (Restifo et al., 2012; Le et al., 2015; Yarchoan et al., 2017; Hilmi et al., 2018; Galon and Bruni, 2019; Duan et al., 2020). To this end, novel immuno-therapeutics which elicit broad anti-tumor T cell responses independent of TCR specificity, such as T-cell bispecific (TCB) antibodies, have shown promise in preclinical testing in immune excluded cancer models (Bacac et al., 2016a; Ishiguro et al., 2017; Bacac et al., 2018; Gedeon et al., 2018; Crawford et al., 2019; Rader, 2020).

In addition to T cell infiltration, intercellular interactions directly within the tumor tissues have been linked with positive response outcomes (Böttcher et al., 2018; Cremasco et al., 2021; Ozga et al., 2021). In particular, interferon gamma (IFN□) produced by CD8 T cells in response to checkpoint blockade, as well as with TCB immunotherapy, has been shown to enhance maturation of intra-tumoral DCs, leading to increased production of chemokine CXC ligand (CXCL)9, CXCL10, and interleukin-12 (IL-12), which in turn promote amplified recruitment, proliferation and differentiation of CD8 T cells (Garris et al., 2018; Chow et al., 2019; Cremasco et al., 2021). Such positive feedback between DCs and CD8 T cells requires direct cell-cell crosstalk, suggesting the need for extensive communication between innate and adaptive immune cells within tumors. Moreover, recent studies have also demonstrated that a subpopulation of CD8 T cells (resource CD8 T cells) which co-express TCF1 and PD1 are not functionally exhausted, and possess a unique capacity for enhanced proliferation and generation of terminally differentiated effector CD8 T cells in response to checkpoint blockade therapy (Im et al., 2016; Wu et al., 2016; Utzschneider et al., 2016; Sade-Feldman et al., 2018; Miller et al., 2019; Siddiqui et al., 2019; Chen et al., 2019; Utzschneider et al., 2020; Krishna et al., 2020). Spatial mapping of these resource CD8 T cells revealed enriched localization near blood vessels in mouse melanomas, and in close proximity to aggregates of antigen presenting cells within vascularized regions of kidney cancers (Jansen et al., 2019; Siddiqui et al., 2019). This suggests the existence of niche-like immune microenvironments within tumors which are likely to be involved in promoting the generation of responses to immunotherapies. As a similar concept, presence of immune-rich tertiary lymphoid structures within tumors have also been linked to disease outcome (Martinet et al., 2011; Wirsing et al., 2014; Hiraoka et al., 2015; Joshi et al., 2015; Finkin et al., 2015; Lee et al., 2016; Sautès-Fridman et al., 2019). Together, these studies suggest that immune cell organization in tumors is critical for effective tumor control with therapy. Nevertheless, the heterogeneous nature of the tumor and the microenvironments it encompasses remains poorly studied, primarily due to the general paucity of tools to study the spatial organization of phenotypically complex cells within irregularly structured tissues.

Here, we utilized multi-parameter confocal imaging coupled with advanced spatial analysis using histocytometry and CytoMAP (Gerner et al., 2012; Stoltzfus et al., 2020a) to examine the complexity of immune cell organization within partially and poorly infiltrated tumors during immunotherapy. To this end, mice bearing MC38 colon carcinomas or KPC pancreatic adenocarcinomas engineered to express human carcinoembryonic antigen (CEA) were treated with CEA-TCB murine surrogate antibody and / or with anti-PD-L1 checkpoint inhibitor, both of which can synergize to promote enhanced CD8 T cell immunity (Bacac et al., 2016a, 2016b; Sam et al., 2020; Cremasco et al., 2021). As expected, CEA-TCB monotherapy and CEA-TCB plus anti-PD-L1 combination therapy led to increased CD8 T cell numbers and decreased tumor burden, indicating control of tumor growth by the immune system (Bacac et al., 2016a; Sam et al., 2020). However, even in these inflammatory settings most CD8 T cells, and particularly the non-exhausted TCF1^+^PD1^+^ resource T cells, were excluded from the active CEA^+^ tumor nest regions. Instead, they were localized in close association with DCs and activated macrophages directly along the perivascular edge of intra-tumoral blood vessels. These perivascular immune aggregates (perivascular immune niches) were detected in untreated tumors and markedly increased in abundance during therapy, indicating active remodeling of the TME during inflammation. Moreover, the relative abundance of these immune-rich microenvironments directly mirrored response efficacy, suggesting their involvement in anti-tumor immunity after therapy. Thus, our studies provide a framework for the application of advanced spatial analysis in studying TME complexity and decoding responses to therapy, as well as reveal that the perivascular immune niche is a microenvironment subtype within tumors with likely involvement in the generation of productive immune responses after therapy.

## Results

### Monotherapy with CEA-TCB and combination of CEA-TCB with anti PD-L1 controls tumor progression in MC38-CEA tumors

To study the composition and spatial patterning of immune cells within tumor tissues in the absence or presence of immunotherapy, we first utilized the MC38 murine colorectal carcinoma model, which has been previously shown to exhibit moderate CD8 T cell responses to checkpoint blockade therapy. To this end, MC38 cancer cells were engineered to express human CEA antigen (MC38-CEA) and inoculated s.c. into CEA transgenic mice, thus mimicking endogenous CEA expression as a tumor-associated antigen. When tumors reached 100-300mm^3^ in volume, animals were randomized into the following treatment groups: one group received the murine surrogate CEA-targeted T cell bi-specific antibody (CEA-TCB), which simultaneously binds to the CEA protein on cancer cells and CD3 on T cells and elicits a T cell mediated attack on CEA-expressing tumors, independent of T cell receptor specificity (Bacac et al., 2016a; Sam et al., 2020); one group was treated with the checkpoint inhibitor, anti-PD-L1 (aPD-L1), and a third group was treated with the combination of CEA-TCB and aPD-L1 antibody. The last group received only vehicle control. As expected, monotherapy with CEA-TCB or aPD-L1 alone resulted in partial tumor control with substantial response variability across individual animals, while combination therapy with CEA-TCB plus aPD-L1 demonstrated enhanced efficacy across the cohort (Fig 1A) (Sam et al., 2020). Evaluation of T cell infiltration at the study endpoint by flow cytometry demonstrated significantly increased frequencies of PD1^+^ and Ki-67^+^ CD8 T cells in CEA-TCB and CEA-TCB plus aPD-L1 therapy groups, suggesting induction of potent CD8 T cell responses (Fig 1B). In contrast, aPD-L1 treatment alone failed to produce a similar magnitude response. Together, these studies confirmed published observations that the CEA-TCB immunotherapy markedly enhances anti-tumor responses by CD8 T cells, and these are further amplified by combination therapy with aPD-L1 (Sam et al., 2020).

**Figure 1:**
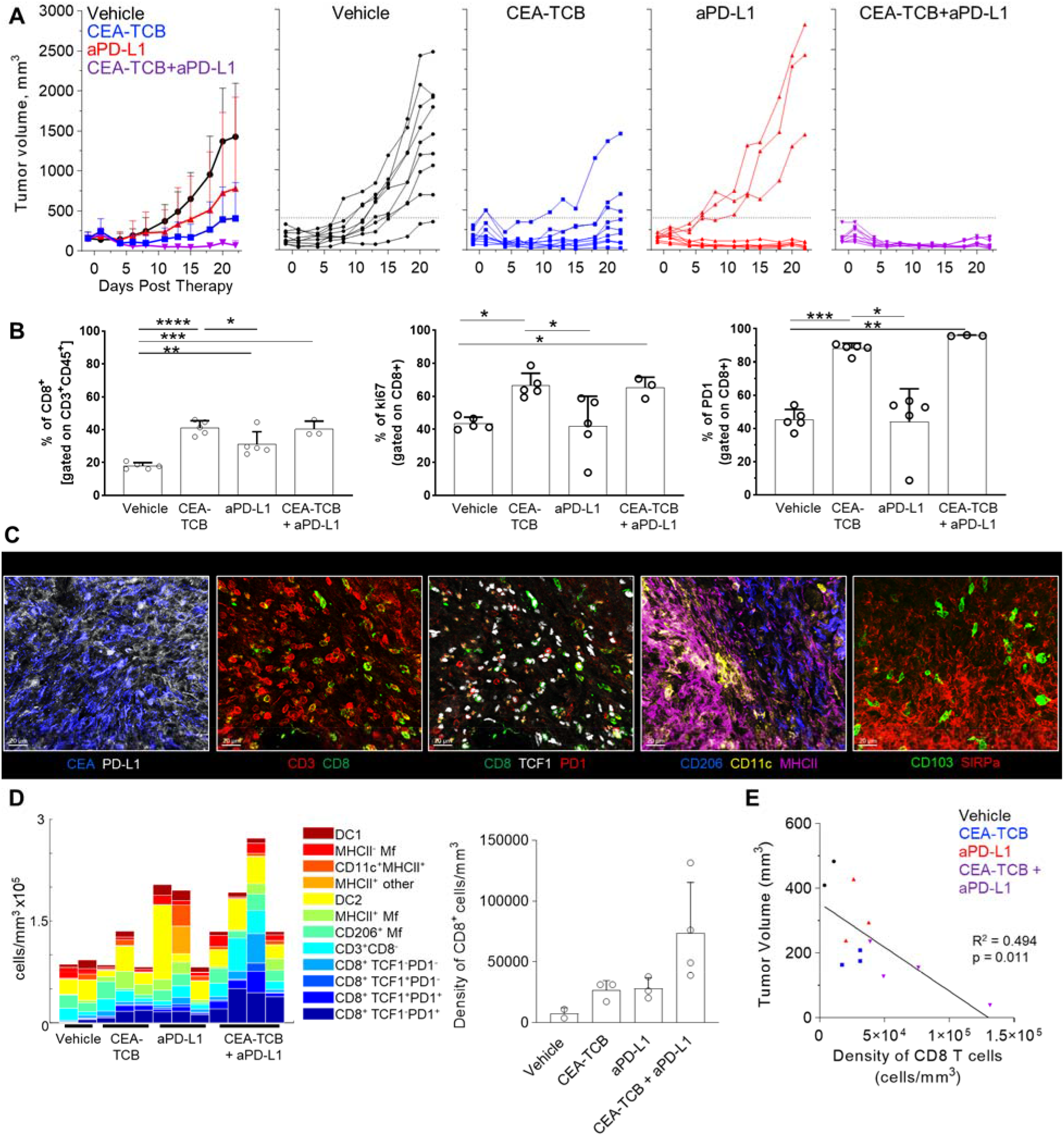
Efficacy of CEA-TCB and anti PDL-1 therapeutic interventions in MC38-CEA tumors. MC38-CEA tumor-bearing mice were treated with the indicated immunotherapies when tumors ranged ~100-300mm^3^ in volume. Treatment regimens were continued in 3d intervals for CEA-TCB and combination, and 7d intervals for aPD-L1. **(A)** Average tumor volume for each group (left), tumor volumes for each sample and treatment (right) after start of treatment. Pooled data from 2 independent experiments (n= 9/group). **(B)** Frequency of CD8 T cell among all T cells (left), the percentage of Ki-67^+^ (center) and PD-1^+^ CD8 T cells (right) in each treatment group as determined by flow cytometry at study end point (22d post start of treatment, n=5/group). **(C)** Multiplex confocal images highlighting markers used to phenotype immune cells in MC38-CEA tumors. Scale bars are 50 μm. **(D)** Average density of immune cell populations for individual tumor samples (columns) per imaged volume (left) and CD8 T cell density across different groups (right), as identified via histocytometry. **(E)** Correlation between the tumor volume and density of CD8 T cells identified by histocytometry. Tumor growth curves and harvest points for data in C-E are displayed in Fig S1A. n=2 for control, n=3 for CEA-TCB and aPDL1 and n=4 for CEA-TCB+aPDL1 group. Data points represent individual tumor samples. Bar graphs show mean, and error bars represent standard deviation (SD). *=p< 0.01, **= p<0.01, ***=p<0.001, ****= p<0.0001.

To investigate the spatial organization of immune cells during early therapy-induced regression timepoints, we carried out multi-parameter confocal imaging of tumor tissues resected four days after initiation of therapy (Fig S1A, red lines indicate imaged samples). Samples were imaged using a 13-plex microscopy panel (Fig 1C, Table S1, S3), and imaged tissues were analyzed using histocytometry and CytoMAP, with identification of twelve major lymphoid and myeloid immune cell types, as well as of cancer-derived CEA signals. Identified myeloid subsets included CD11c^+^MHCII^+^ dendritic cells (DCs), further composed of CD103^+^ DC1 and SIRPα^+^ DC2, CD11c^−^ CD206^+^ macrophages (Mfs), as well as CD11c^−^ CD206^−^ SIRPa^+^ Mfs, which were further sub-gated based on major histocompatibility complex II (MHCII) expression. Identified lymphocyte populations included CD8 T cells, which were further stratified based on TCF1 and PD1 expression, as well as CD3^+^ CD8^−^ cells (putative CD4 T cells). PD-L1 expression in imaged tissues was also assessed (Figs 1C, S1B).

In accordance with the flow cytometry data (Fig 1B), combination therapy with CEA-TCB plus aPD-L1 markedly enhanced CD8 T cell infiltration, and this was primarily associated with the expansion of TCF1^−^PD1^+^ CD8 T cells (Figs 1D, S1C). While these T cells likely represent a complex mixture of effector and exhausted populations, due to lack of additional markers to further discriminate these subsets, the TCF1^−^ PD1^+^ population will be referred to as effector CD8 T cells (Blackburn et al., 2009; Fourcade et al., 2010; Matsuzaki et al., 2010; Im et al., 2016; Hashimoto et al., 2018; Sade-Feldman et al., 2018; Miller et al., 2019; Siddiqui et al., 2019; Jiang et al., 2021). Less dramatic increases with therapy were seen for the TCF1^+^PD1^+^ CD8 T cells (Figs 1D, S1C), a recently identified resource T cell population which undergoes proliferation in response to immunotherapy and gives rise to downstream terminal effector and exhausted cells (Im et al., 2016; Sade-Feldman et al., 2018; Jadhav et al., 2019; Siddiqui et al., 2019; Chen et al., 2019; Krishna et al., 2020). CEA-TCB or aPD-L1 monotherapy groups also demonstrated partial expansion of both CD8 T cell subsets, but were less efficacious than the combination treatment. Importantly, consistent with past studies, increased density of CD8 T cells was negatively correlated with tumor volume, indicating control of disease progression by this immune cell type (Fig 1E) (Hanson et al., 2000; Pagès et al., 2005; Galon et al., 2006; Mahmoud et al., 2011; Bacac et al., 2016a; Sam et al., 2020). With regard to myeloid cells, we noted substantial prevalence of DC2s and Mf populations across all conditions, with partial increases in MHCII^+^ activated Mfs after CEA-TCB mono and combination treatments (Fig 1D). In contrast, relatively minor representation of DC1s was seen across all samples and treatments.

### Quantitative analysis of the MC38-CEA tumor microenvironment

We next examined the organization of CD8 T cells across the different experimental conditions. Increased CD8 T cell abundance was observed for all treatments as compared to control samples with substantially greater density after CEA-TCB plus aPD-L1 treatment (Fig 2A). Complex heterogeneous patterns of T cell infiltration were also observed across all treatment groups. To explore how the different T cell subsets were distributed throughout the tumor, we first quantified the degree of cellular infiltration into the CEA-expressing tumor regions using a simple distance-based approach. For this, MC38-CEA tumors were characterized based on the distribution of cancer cell derived CEA signal (CEA spot objects), as well as of CD206^+^ Mfs, which were found closely associated with the outer capsular edges of the tumors (Fig S2A). Using these parameters, we computationally defined the CEA-expressing tumor nest with CytoMAP (Fig 2B) and calculated the distance of different CD8 T cell subsets to this tumor boundary (Fig 2C, Fig S2B). As expected, this analysis demonstrated preferential localization of CD206^+^ Mfs externally to the CEA^+^ tumor boundary, as well as the internal localization of MC38-CEA cells (Fig 2C, Fig S2B). This also revealed that the two CD8 T cell subsets were differentially distributed within the tissues. The TCF1^+^PD1^+^ resource CD8 T cells were predominantly, but not exclusively, located closer to the tumor border as compared to the TCF1^−^ PD1^+^ effector CD8 T cells, which were located further away from the border, suggesting deeper infiltration (Fig 2C, Fig S2B). These differences were observed in all treatment groups, albeit the degree and depth of infiltration varied across experimental conditions and individual samples (Fig 2C, S2B). Together, these findings indicate that CD8 T cells increase in cellularity and at least partially infiltrate the tumors after immunotherapy, as well as that the TCF1^+^PD1^+^ progenitor and TCF1^−^PD1^+^ effector CD8 T cells have non-equivalent spatial distribution properties within the tumor tissues.

**Figure 2:**
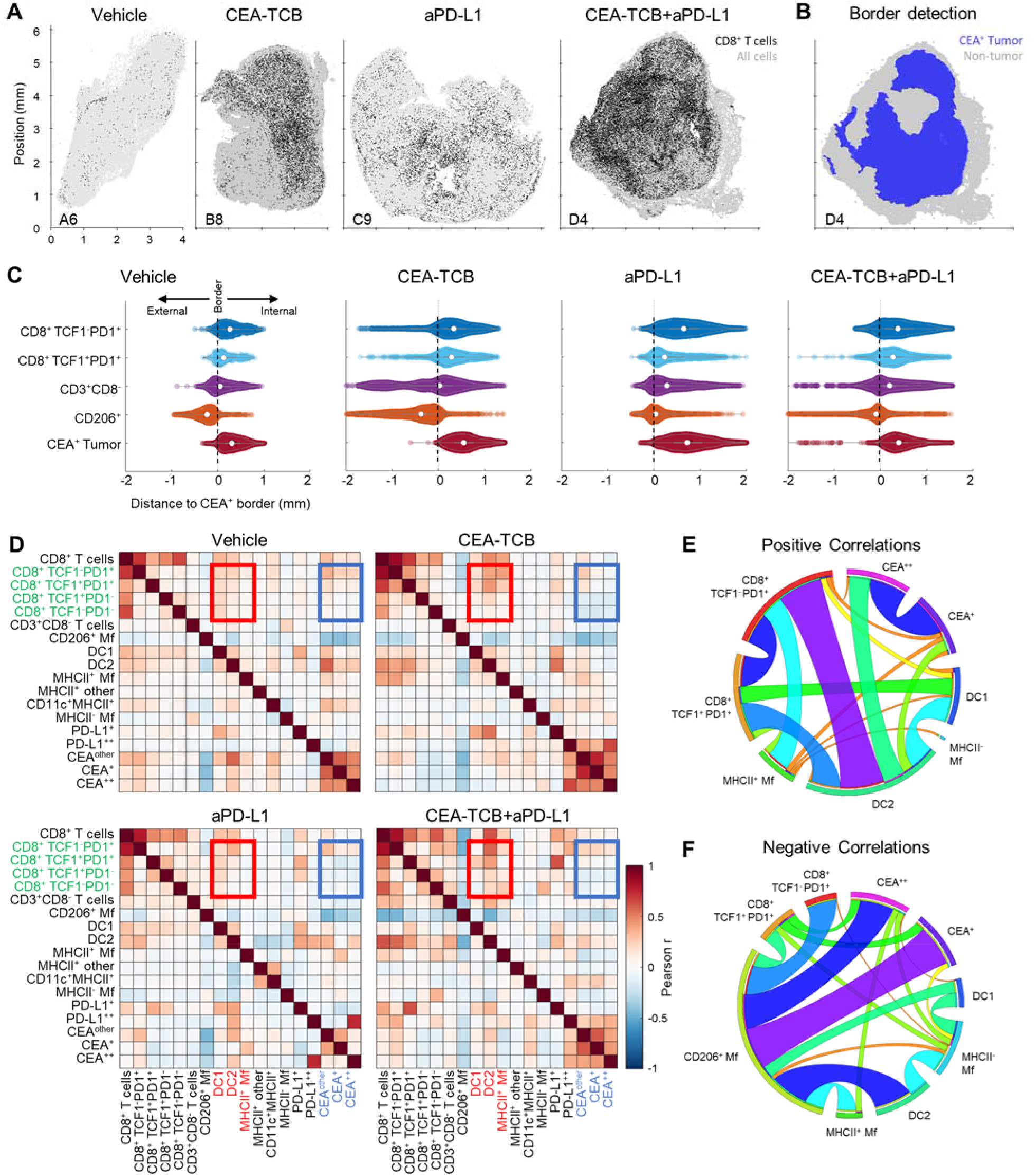
Histocytometry and CytoMAP based quantification of the MC38-CEA tumor microenvironment. **(A)** Positional plots of CD8 T cells (black) overlaid over all imported cell objects (gray) in the indicated tumor samples. **(B)** CEA^+^ tumor regions were identified using CEA spot clustering and visualized (blue), with additional overlay of all other objects (grey). **(C)** The distances of select cell populations to the border of the CEA^+^ tumor region is shown in Fig. 2B from combined samples of each treatment group. Negative distances correspond to positions outside the border of the tumor region and positive distances correspond to positions inside the tumor region. **(D)** Heatmaps displaying the Pearson’s correlation between number of cells per neighborhood for each cell population pair for each treatment group. **(E)** Circos plots of the scaled and rounded positive Pearson correlation between number of cells per neighborhood across all samples for the indicated cell types. Plots were generated using www.circos.ca. **(F)** Circos plots of the scaled and rounded negative Pearson correlation between the number of cells per neighborhood across all samples for the indicated cell types. Colors were auto-generated. All the above plots were generated as described in Fig S1A; n=2 for control, n=3 for CEA-TCB and aPDL1 and n=4 for CEA-TCB+aPD-L1 group.

To further interrogate cellular patterning, we used CytoMAP to quantify the spatial correlations between the different immune cell subsets and across conditions (Fig 2D-F). This analysis identifies cell types which preferentially localize near each other (positive correlation), or conversely avoid one another (negative correlation) within the tissues. The Pearson correlation coefficient was calculated for the number of cells of each population pair within 50 μm raster-scanned spatial neighborhoods across all samples and visualized (Fig 2D-F). This revealed that in all conditions T cells were positively correlated with one another, suggesting that T cells generally tend to be colocalized (Fig 2D, 2E). In contrast, most T cells, and in particular the resource TCF1^+^PD1^+^ CD8 T cell population, displayed either a neutral or negative correlation with CEA^+^ spots, suggesting a general exclusion from the tumor nest areas (Fig 2D, 2F). The only T cells that displayed a positive correlation with CEA^+^ spots were the effector TCF1^−^PD1^+^ CD8 T cells, indicating partial infiltration of the deeper tumor nest regions by this population and corroborating results obtained from the distance-based analysis (Fig 2C, S2B). Furthermore, both CD8 T cell subsets were positively correlated with DCs, and to a lesser degree with activated MHCII^+^ Mfs, but negatively correlated with non-activated SIRPa^+^CD11c^−^MHCII^−^ Mfs (Fig 2D-F). Of note, while positive associations with DC1s were observed, the DC2 subset was substantially more abundant in all examined samples (Fig 1D). In addition, T cells were positively correlated with intermediate PD-L1 expression on cells in surrounding neighborhoods (Fig 2D). In contrast, high PD-L1 signal was correlated with the location of CEA^+^ spot objects representing cancer cells, and this correlation was further enhanced after CEA-TCB mono and combo therapy, consistent with immune-mediated modulation of this molecule (Bacac et al., 2016a; Sam et al., 2020).

To globally investigate the organization of all immune cells within the MC38-CEA tumors, we next performed neighborhood clustering and region analysis. For this, raster-scanned spatial neighborhoods were clustered using a self-organizing map and regions of similar cellular representation were manually concatenated and annotated (Fig 3A). This revealed eleven distinct regions (R1-R11) with varying abundance of distinct lymphoid, myeloid and cancer cells, including: MHCII^−^ or CD206^+^ Mf enriched (R2-R3), T cell dense (R4), TCF1^+^ CD8 T cell and DC2 enriched (R5), general T cell – DC2 rich (R6), T cell – MHCII^+^ activated Mf (R7), myeloid-rich with cancer cells (R8-9), as well as CEA^+^ tumor and tumor nest regions (R10-11) which were devoid of T cells (Fig 3A). Neighborhoods belonging to the identified regions were visually verified for the appropriate cellular composition (Fig S3A).

**Figure 3:**
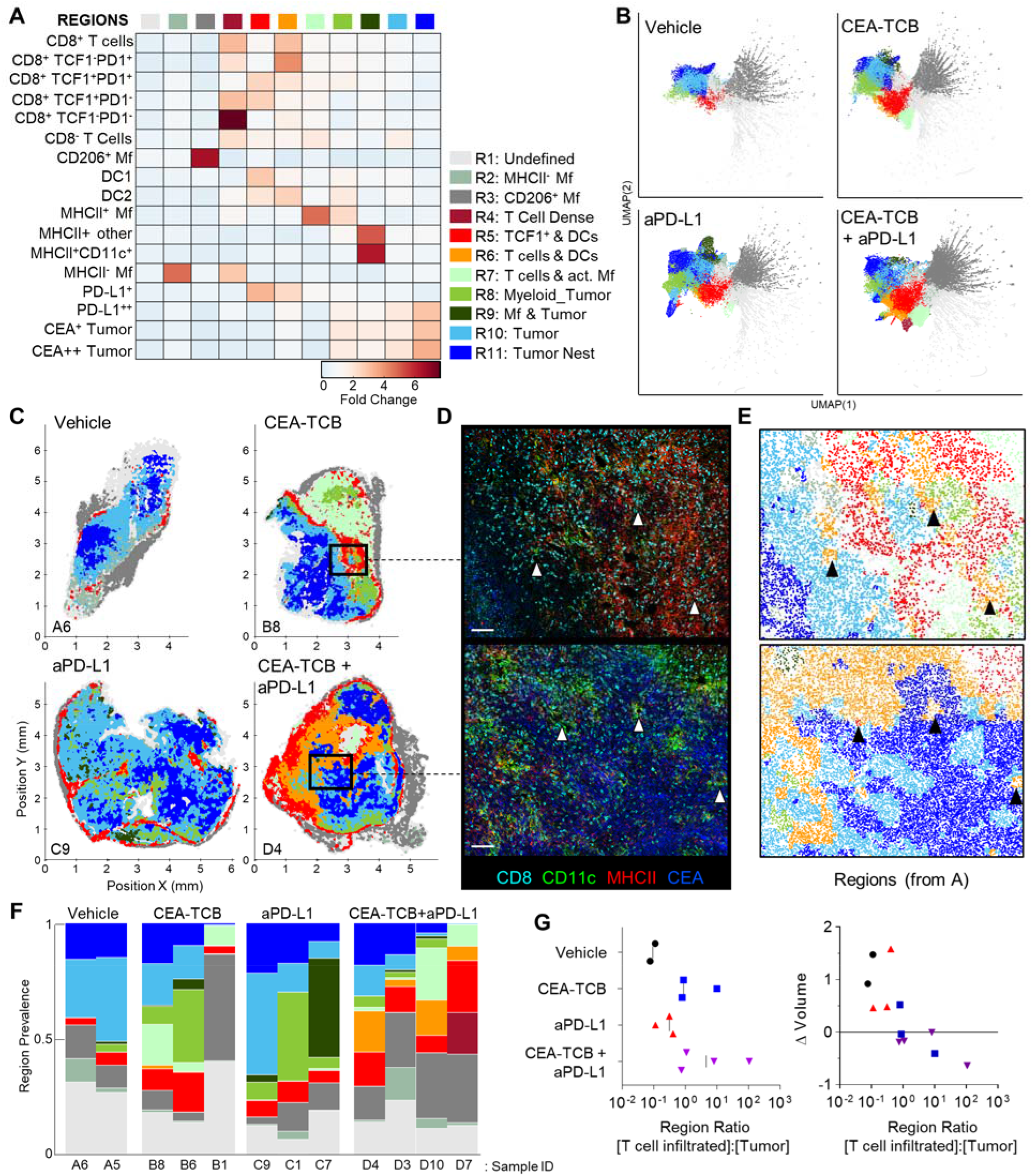
CytoMAP based regional analysis of the MC38-CEA tumor microenvironment. **(A)** Spatial neighborhoods were clustered into regions and plotted on a region heatmap. Plot displays fold change of object density per neighborhood within the indicated regions across all samples. **(B)** UMAP plots of the color-coded neighborhoods from each treatment group displaying the heterogeneity in the tissue regions. **(C)** Plots of the positions of all neighborhoods for select samples from each treatment group, color coded by region classification defined in Fig. 3A. **(D)** Multiplexed confocal regions of interest boxed in Fig. 3C. Arrows highlight select niches enriched in T cell and DCs. Scale bar represents 100 μm. **(E)** Plots showing the individual CytoMAP objects for the regions of interest shown in Fig. 3D, color-coded by region classification defined in Fig. 3A. **(F)** Region prevalence for each sample (columns) within the treatment groups. **(G)** Ratio of immune infiltrated regions to immune excluded regions (left). Region ratio plotted versus fold change in tumor volume after the initiation of therapy. All the above plots were generated as described in Fig S1A; n=2 for control, n=3 for CEA-TCB and aPDL1 and n=4 for CEA-TCB+aPD-L1 group.

Unsupervised dimensionality reduction of the neighborhoods based on cellular composition was also performed using UMAP or t-SNE (Fig 3B, S3B) (van der Maaten and Hinton, 2008; McInnes et al., 2018). This visualization demonstrated clustering of similar neighborhoods, with different region types identified in Figure 3A (color-coded) also showing close alignment with the clusters. Moreover, this analysis demonstrated major changes in region representation based on condition, with marked enhancement of multiple T cell & DC enriched regions (R4-R6) after immunotherapy (Figure 3B). Given that inter-cluster distances on the UMAP plots directly reflect the degree of similarity between the clusters, with the connections between clusters also representing likely transitions between the regions, we used these plots to study general region organization within the tumors. This indicated that the R3 CD206^+^ Mf regions (dark grey) were most separated from the rest of the neighborhoods (Fig 3B), consistent with the segregated capsular localization of this myeloid cell type (Fig S2A). The R3 CD206^+^ Mf neighborhoods were connected to the rest of the neighborhoods via the R5 region (red), enriched in TCF1^+^PD1^+^ resource and TCF1^−^PD1^+^ effector CD8 T cells as well as in DCs (Fig 3B). The R5 region was in turn connected to the tumor rich regions (R8-R11). In treated samples, the R5 region was also connected to additional T cell rich regions with stronger representation of DC2 and activated MHCII^+^ Mfs (R4, R6, R7), albeit these were separated from CEA^+^ tumor regions within the UMAP space (Fig 3B). These transitions indicate likely structural organization of the regions with respect to one another, with the T cell and DC rich R5 region serving as a bridge linking the outer CD206^+^ Mf capsular region with the rest of the tumor, and with likely segregation of additional T cell rich neighborhoods from the tumor nest.

Direct visualization of region distribution highlighted complex spatial patterning within individual tissues and across conditions (Figure 3C-E). Some noted associations regarding CD206^+^ Mfs lining the capsular border and CEA^+^ spots defining tumor nest regions were confirmed across samples, while abundant T cell rich regions could be observed surrounding the CEA^+^ tumor nest regions in the CEA-TCB plus aPD-L1 samples (Fig 3C). The R5 region was primarily located in the periphery of the samples, in close association with R3 CD206^+^ Mf region, while the T cell rich (R4-R7) and CEA+ tumor nest regions (R8-R11) appeared segregated from one another (Fig 3C). These data indicate formation of discrete foci of immune reactivity and heterogeneous distribution of distinct TMEs across the tissues. Moreover, while the simpler distance and spatial correlation analyses indicated partial infiltration of the tumor bed by the TCF1^−^PD1^+^ effector CD8 T cells, the global region-based analysis also demonstrated that, in general, T cell rich and CEA^+^ tumor nest regions are spatially segregated from one another, and that even after immunotherapy, CEA^+^ tumor nests are relatively devoid of T cells.

We next quantified region prevalence across different conditions (Fig 3F). Each treatment group appeared to alter region representation in unique ways, with the combination group displaying the most dramatic shifts, with markedly increased abundance of the (R5) TCF1^+^ & DC rich and (R6) T cell & DC rich regions. In contrast, aPD-L1 monotherapy was associated with increased representation of Mf or myeloid rich tumor regions (R8, R9), and a lower abundance of T cell infiltrated regions.

Substantial intra-group variability in region prevalence was also noted and we explored whether this heterogeneity was related to disease progression of individual animals. Since CEA-TCB mono- and combination therapies were strongly associated with increased representation of several T cell rich regions (Fig 3F), we calculated the total sum of these T cell dense regions using CytoMAP. This sum was negatively correlated with the fold change in tumor volume after initiation of treatment, supporting the notion that enhanced T cell numbers and function after immunotherapy can promote tumor regression (Fig S3C). In contrast, untreated or aPD-L1 only treated samples had higher representation of the T cell excluded, CEA^+^ tumor regions, and the sum of these regions was associated with increased fold change in tumor volume, suggesting disease progression (Fig S3D). These relationships were further explored by calculating the ratio of T cell rich regions to CEA^+^ tumor regions. As expected, this calculated ratio was higher in CEA-TCB and CEA-TCB plus aPD-L1 combination groups, and importantly, was negatively associated with fold change in tumor volume (Fig 3G). Of note, even with the highly variable region representation among the individual tumors (Fig 3F), the calculated region ratio clearly aligned all samples along the same trajectory (Fig 3G). This indicates that even with extensive heterogeneity across samples and conditions, the relative representation of T cell infiltrated vs. tumor nest regions may serve as an accurate reflection of ongoing immune responses to therapy.

### Perivascular immune niches are a major inflammatory microenvironment in MC38-CEA tumors

Our spatial correlation and neighborhood clustering analyses revealed a strong relationship among CD8 T cells, DCs, and activated MHCII^+^ Mfs (Fig 2D, 3A), suggesting an intimate association between these cell types. Indeed, closer visualization of tumor cross-sections confirmed robust presence of T cells in regions heavily populated by CD11c^+^ DCs and MHCII^+^CD11c^−^ activated Mfs (Fig 3D). These innate cells also appeared to be aggregated in intricate formations, generating structures akin to corridors or small islands segregated away from CEA^+^ tumor nest regions, and were localized around smaller unstained structures which appeared similar in morphology to blood vessels. To study the relationships between immune cells and tumor blood vessels, we designed a new imaging panel which incorporated CD31 vascular endothelium staining. Indeed, visual inspection of the imaged tissues revealed remarkable clustering of DCs and activated Mfs directly along the perivascular cuff of intra-tumoral blood vessels (Fig 4A). Infiltrating CD8 T cells, both TCF1^+^PD1^+^ and TCF1^−^PD1^+^, were also enriched in these perivascular regions. Presence of this perivascular immune microenvironment (perivascular niche) was observed in untreated samples, but was greatly increased in abundance after immunotherapy, especially in the CEA-TCB treatment groups (Fig. 4A). To quantify these relationships, the spatial distribution of various myeloid and lymphoid cell objects, as well as of CD31 blood vessel objects was analyzed using CytoMAP. Neighborhood clustering revealed that as before, most CD8 T cells were enriched in similar regions as DCs (R2) or activated Mfs (R3) (Fig 4B). Blood vessels were also highly enriched in these regions, supporting the perivascular localization of these immune populations. Region prevalence and dimensionality reduction analyses demonstrated presence of the DC – T cell perivascular microenvironment (R2) in untreated samples, as well as marked increases after CEA-TCB combination treatment (Figs 4C, 4D, S4A), being in close alignment with previous analyses (Fig 3B, 3F). Furthermore, the spatial localization of these perivascular immune niches was observed to be predominantly restricted around the outer edge of the tumor nest or along the capsular border, indicating general segregation from the internal tumor nest compartment (Fig 4D).

**Figure 4:**
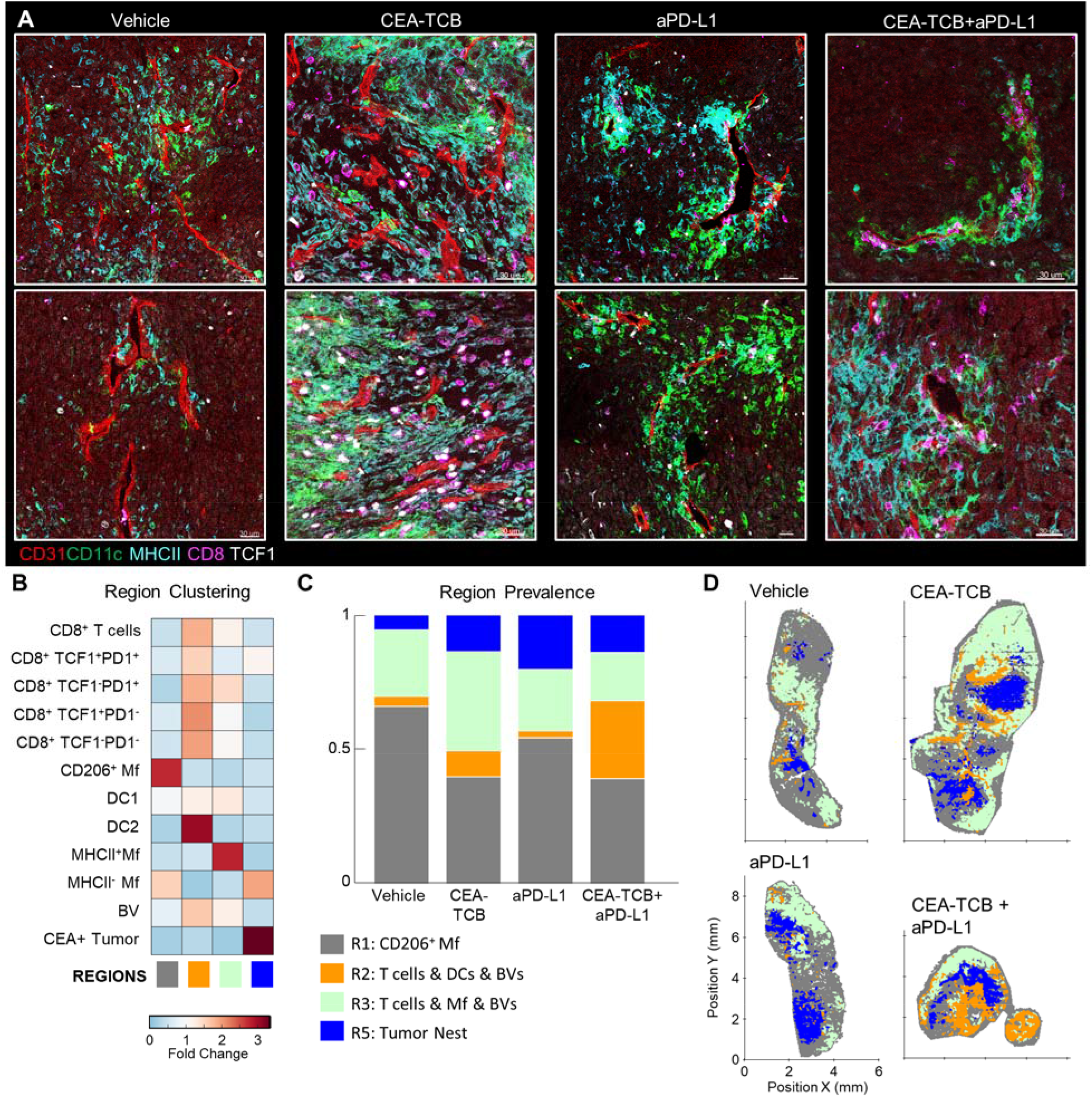
Perivascular immune niches in MC38-CEA tumors. **(A)** Multiplex confocal images showing the association between TCF1^+/-^ CD8 T cells and DCs (CD11c+ MHCII+) with CD31+ blood vessels (BVs) across the different treatment conditions. **(B)** Neighborhoods (50μm) were clustered into regions and plotted on a region heatmap. Plot displays fold change of object density per neighborhood across all samples for each region. **(C)** Region prevalence in each sample. **(D)** Positional plots of all neighborhoods for selected samples, color-coded by region classification defined in Fig. 4B.

To further explore the associations of different immune populations with tumor blood vessels, we again performed spatial correlation analysis. This confirmed the observed relationships with strong positive correlation of T cells, DC2s, and activated Mfs with blood vessels, as well as general exclusion of CEA^+^ cancer spot objects (Fig S4B). We also calculated the distances of immune cells to the nearest blood vessels. This revealed highly proximal positioning of DC2 near blood vessels as compared to the non-activated MHCII^−^ Mfs (Fig S4C, left). Differences in vessel proximity between TCF1^+^PD1^+^ resource and TCF1^−^PD1^+^ effector CD8 T cells were also noted S4C, middle). In untreated samples, both subsets were generally located highly proximal (<25μm) to blood vessels. However, immunotherapy increased the distance of effector, but not resource, CD8 T cells to blood vessels, indicating partial infiltration of the deeper tumor regions by this population. Consistent with this, effector T cells were found in closer proximity to CEA^+^ spot objects compared to resource CD8 T cells, and this distance was further decreased with combination therapy (Fig S4A, right). Thus, the combination of region- and distance-based analyses demonstrate distinct but complementary information, that most CD8 T cells are localized in highly vascularized DC-rich microenvironments, and that during initiation of responses to immunotherapy, expanded effector, but not resource, CD8 T cells can infiltrate the tumor bed.

### Perivascular immune niches in additional tumor models

We next examined whether similar perivascular immune microenvironments could be observed in additional tumor models. We first visualized CT26 colorectal carcinomas and B16.F10 melanomas and detected strong spatial associations between activated DCs, T cells and blood vessels (Figs S4D, S4E). We then turned to the KPC pancreatic ductal adenocarcinoma model, which is characterized as being highly aggressive, poorly infiltrated by T cells, and resistant to checkpoint blockade therapy. KPC cells were engineered to express CEA antigen and inoculated into CEA transgenic mice. As above, when the tumors reached 100-300mm^3^, animals either received vehicle control injections, or were treated with the CEA-TCB and aPD-L1 mono- or combination immunotherapies. Treatment with CEA-TCB or CEA-TCB plus aPD-L1, but not aPD-L1 alone, elicited modest changes in tumor growth suggesting partial immune mediated protection (Fig 5A). Consistent with this, CEA-TCB mono and combination treatments were associated with enhanced numbers of CD8 T cells, dominantly driven by the expansion of TCF1^−^ PD1^+^ effector T cells, as well as with moderate increases in activated myeloid cells in the combination treatment group (Fig 5B-D). While substantial heterogeneity in cellular abundance between samples was noted, in general, increased density of CD8 T cells after CEA-TCB therapy was negatively correlated with tumor volume, indicating partial control of tumor growth by the activated T cells (Fig 5E).

**Figure 5:**
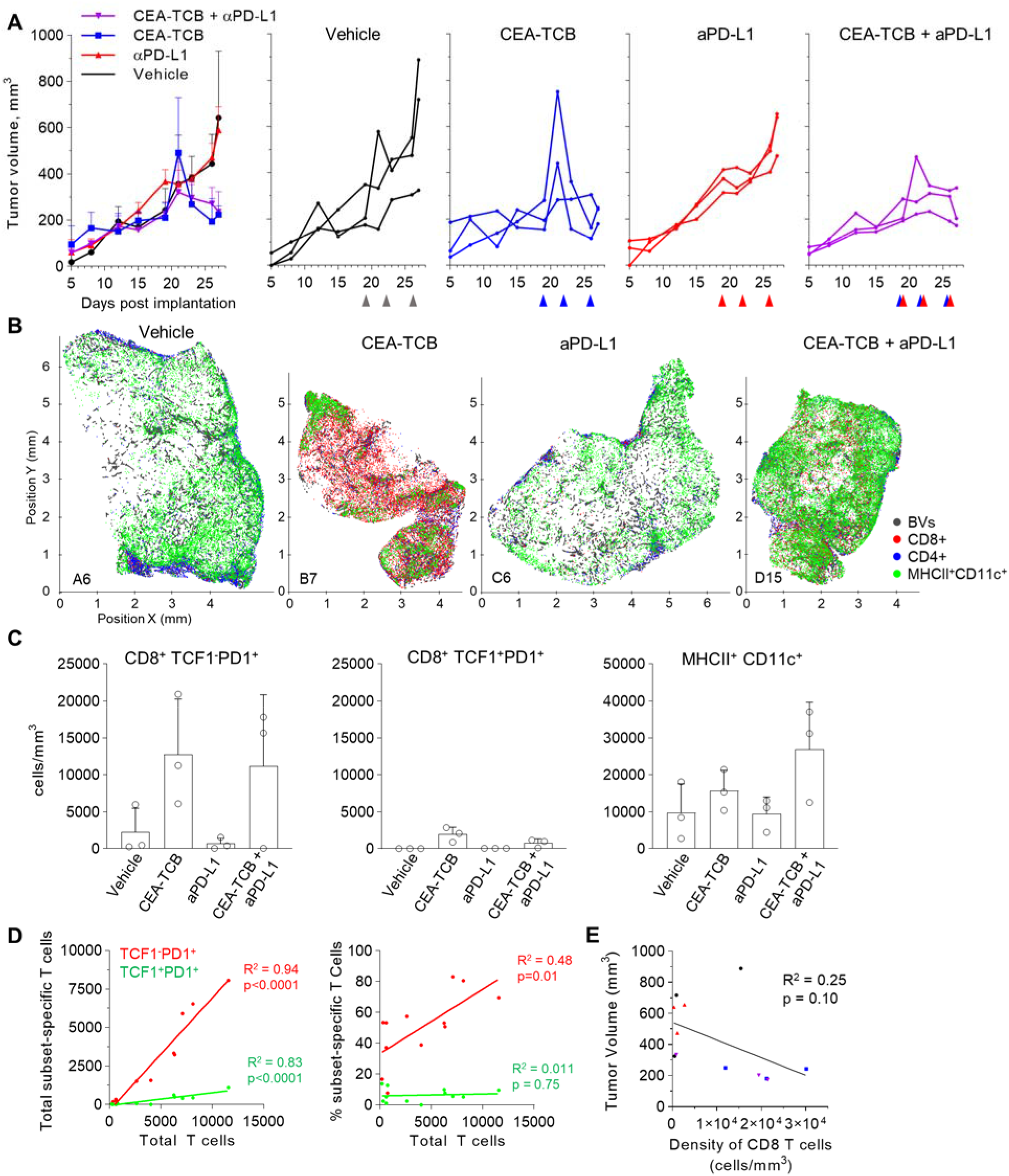
Efficacy of CEA-TCB and anti PDL-1 therapeutic interventions in the KPC-CEA tumor model. **(A)** KPC-CEA tumor-bearing mice treated with the indicated immunotherapies starting 19 days after implantation. Treatments were continued in 3d intervals as shown by arrows. Average tumor volume (left), and volume of individual samples (right) from each treatment group are shown. Data from one independent experiment (n= 3/group). **(B)** Positional plots of color-coded objects defined by histocytometry for select samples from each treatment group. **(C)** Density of CD8^+^TCF1^−^PD1^+^ (left), CD8^+^TCF1^+^PD1^+^ (center), or MHCII^+^CD11c^+^ (right) cells by treatment group as identified by histocytometry. **(D)** Correlation between the total number of CD8 T cells and either the total number (left) or percent (right) of the indicated T cell subsets. **(E)** Correlation between the tumor volume and the density of CD8 T cells as identified by histocytometry. Data points represent individual samples. Bar graphs show mean, and error bars represent SD.

We further analyzed the organization of immune cells within the KPC-CEA tumors using CytoMAP. As before, 50μm raster-scanned spatial neighborhoods across all imaged samples were clustered, with subsequent manual concatenation and annotation (Fig 6A). This revealed seven distinct region types with varying abundance of lymphoid, myeloid, blood vessels and cancer cells, including: (R2) blood vessel rich, (R3) CD4^+^ T cell and myeloid cell region with blood vessels, (R4) CD4 and CD8 T cell rich region with blood vessels, (R5) highly T cell and DC2 rich region with abundant blood vessels, as well as (R6) Mf rich and (R7) CEA^+^ tumor nest regions (Fig 6A). The clustering of neighborhoods into regions was also visualized with UMAP analysis, which again revealed additional structure to region associations and marked separation of T cell rich vs. CEA^+^ tumor nest neighborhoods (Fig 6B). Based on these data, the R5 region composed of T cells, DC2s and blood vessels appeared highly similar to the perivascular niche identified in the MC38-CEA model (Fig 4A). This was verified by visual inspection of confocal images as well as of CytoMAP annotated cells, revealing close spatial associations of T cells and DCs with intra-tumoral blood vessels (Fig 6C). These positional relationships were also quantified by calculating the distance to nearest blood vessels, or other anatomical and cellular landmarks (Fig S5). DCs were localized in closer proximity to blood vessels compared to non-activated Mfs or cancer cells (Fig S5A). Similarly, the TCF1^+^PD1^+^ resource CD8 T cells were generally more proximal to blood vessels, the tumor border, and DCs as compared to TCF1^−^PD1^+^ effector CD8 T cells, indicating distinct spatial properties for these two CD8 T cell populations (Figs S5A-D). Spatial correlation analysis was also performed, which similarly revealed positive correlation of T cells and DCs with tumor blood vessels, as well as relative exclusion from neighborhoods rich in non-activated MHCII^−^ Mfs and CEA^+^ cancer spot objects across most treatment groups (Figs. S6A-C).

**Figure 6:**
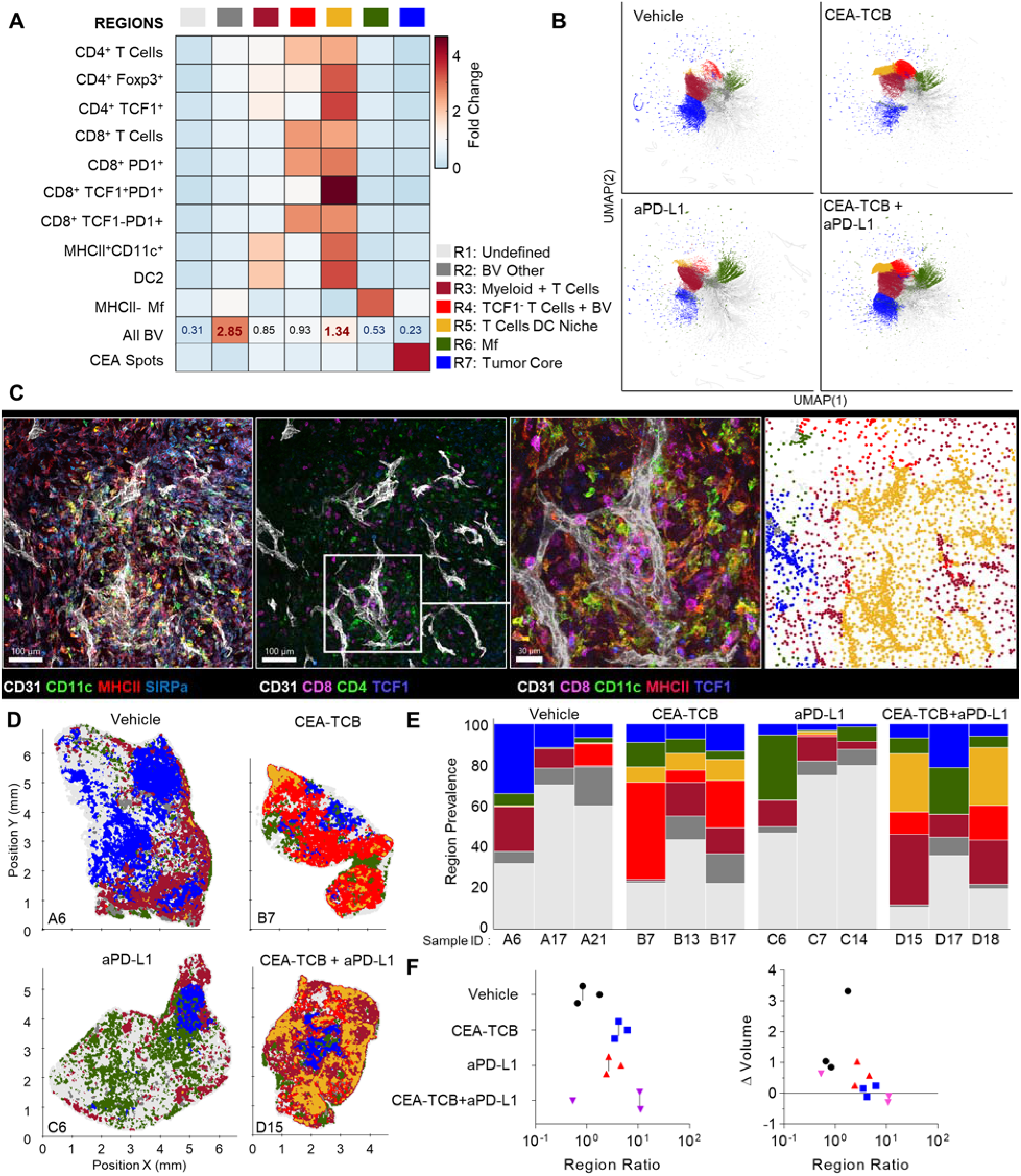
CytoMAP based regional analysis of the KPC-CEA tumor microenvironment. **(A)** Neighborhoods were clustered into regions and plotted on a region heatmap. Plot displays fold change of the object density per neighborhood within each region across all samples. **(B)** UMAP plots of the color-coded neighborhoods from each treatment group showing heterogeneity in the tissue regions. **(C)** Region of interest (ROI) from multiplexed confocal image showing association of the T cells and DCs with BVs. Scale bar represents 100 μm (two left images) and 30 μm (third image). The rightmost plot shows the positions of cells in the same ROI color-coded by the region classification defined in Fig. 6A. **(D)** Positional plots of all neighborhoods for select samples from each treatment group, color-coded by region classification defined in Fig. 6A. **(E)** Region prevalence for each sample (columns) within the four treatment groups. **(F)** Ratio of immune infiltrated regions to immune excluded regions (left). Ratio from F, plotted versus the fold change in tumor volume after initiation of therapy (right). All the above plots were generated from samples collected at d25, n=3 per group.

These data indicated that similar to the MC38-CEA model, KPC-CEA tumors also generate the perivascular immune niche. Furthermore, CEA-TCB mono and combination therapies, but not aPD-L1 treatment alone, markedly increased the representation of this perivascular niche (R5), although as in the MC38-CEA model, substantial variation in region representation for individual samples was noted (Fig 6E). Thus, the perivascular region can expand in size with immunotherapy, reflecting changes in the TME during inflammation and supporting a potential role for these regions in promoting anti-tumor immunity. Indeed, the sum of T cell dense regions (R3-R5) or the ratio of T cell dense to CEA^+^ tumor regions was negatively correlated with fold change in tumor volume post initiation of treatment (Figs 6F, S6A, S6B). Moreover, strikingly similar response patterns were seen across both MC38-CEA and KPC-CEA models, with close alignment of all examined samples along the same general trajectory irrespective of tumor type, or inter- and intra-group heterogeneity (Fig S6C). These data support the premise that quantitative imaging of the TME can reveal fundamental features of immune cell organization within tumors, as well as identify useful spatial biomarkers of immune responses and outcomes after immunotherapy.

## Discussion

Existing dissociation-based technologies, such as multiparameter flow cytometry or single cell RNA sequencing, have revealed a large spectrum of immune cell populations with diverse phenotypic and functional properties, which can infiltrate tumors and influence disease progression (Pagès et al., 2005; Anichini et al., 2010; Jochems and Schlom, 2011; Puram et al., 2017; Azizi et al., 2018). However, understanding how these populations interact and influence one another within tissues inherently requires the use of microscopy. To date, this has been challenging given the general paucity of multiplex imaging and spatial analytics solutions capable of dissecting the organization of phenotypically complex cells within highly heterogenous tissues. Here, we employed high-resolution multiparameter confocal imaging, histocytometry, and computational spatial analysis with CytoMAP to resolve the complexity of the TME in two preclinical murine cancer models, the MC38 colorectal and KPC pancreatic tumors with and without immunotherapy. Our quantitative imaging tools revealed conserved subclasses of microenvironments despite substantial intergroup, sample-to-sample, and intra-tissue variation in cellular patterning. Our analyses demonstrated the existence of perivascular immune niches, which were highly enriched in DCs and resource CD8 T cells as well as other activated cell types. This TME subtype was present in untreated tumors and underwent dramatic expansion after immunotherapy. The relative abundance of immune-rich vs. cancer nest associated microenvironments directly correlated with tumor burden regression in both cancer models, and largely accounted for the heterogeneity in responses of individual animals. These observations support the established notion that activated T cells can exert substantial immune pressure on tumors after immunotherapy, as well as indicate that modular behavior of pre-existing immune microenvironments can set the balance point in anti-tumor responses after therapy.

The finding that CD8 T cells remain largely excluded from the deep tumor nest regions of ‘immunologically cold’ tumors, despite aggressive immunotherapy and substantial CD8 T cell expansion, indicates the presence of potent mechanisms regulating cellular trafficking to and within these tissues. Localization of immune cells in tumors is governed by the distribution of chemokine signals and extracellular matrix components generated by cancer associated fibroblasts, suppressive macrophages, or other cell types, which themselves respond to local tissue cues, such as hypoxia and TGFβ (Chow and Luster, 2014; Joyce and Fearon, 2015; Ozga et al., 2021). The few CD8 T cells that did infiltrate the deeper CEA+ tumor nest regions were enriched in the TCF1^−^PD1^+^ population, while most TCF1^+^ PD1^+^ resource CD8 T cells remained in peripheral regions or within the perivascular immune niche. It is important to re-emphasize that the TCF1^−^PD1^+^ CD8 T cells visualized in this study likely encompass both terminal effector and exhausted populations, but these could not be distinguished due to lack of appropriate markers in our panels. Regardless, these data suggest that different T cell subsets have divergent capabilities for intra-tumoral trafficking, which is highly consistent with reported differences in expression of chemokine receptors and adhesion molecules (Hashimoto et al., 2018; Chen et al., 2019; McLane et al., 2019; Siddiqui et al., 2019; Yao et al., 2021; Seo et al., 2021). Our findings on the distribution of resource T cells are also in direct concordance with two independent reports demonstrating preferential presence of resource T cells in close proximity to blood vessels (Siddiqui et al., 2019) or antigen presenting cells (Jansen et al., 2019) in mouse melanomas and human kidney tumors, respectively. Similar general relationships for the distribution of stem-like resource vs. exhausted effector CD8 T cells were noted in spleens and lymph nodes during chronic LCMV infection (Im et al., 2016, 2020; Duckworth et al., 2021). This indicates a global conservation of divergent spatial trafficking programs for different T cell subsets across conditions, organs, tumor types, as well as species.

It also stands to reason that the distribution of T cell subsets within the tumor is a direct reflection of cellular function. Non-exhausted resource CD8 T cells localize near blood vessels and distal to the deeper tumor regions, while effector T cells can be recruited deeper into the nest, but likely undergo progressive exhaustion with repeated activation or continued exposure to immunosuppressive cues. The close spatial association of resource T cells and DCs near blood vessels may also enhance T cell proliferation in response to tumor antigens presented by the proximal innate subsets, especially following immunotherapy. In turn, this could lead to continued localized seeding of tumors with the generated effector T cells. Inflammatory signals produced by the activated T cells could also lead to further activation of neighboring myeloid and endothelial cells, promoting recruitment of additional resource and effector T cells from the vasculature, which has been observed after both checkpoint blockade and CEA-TCB therapies (Joyce and Fearon, 2015; Spitzer et al., 2017; Huang et al., 2018; Munn and Jain, 2019; Chow et al., 2019; Wu et al., 2020; Zhang et al., 2021; Cremasco et al., 2021; Pauken et al., 2021; Connolly et al., 2021). Additional chemoattraction and activation of T cells already present in the tumor is also likely (Chow et al., 2019). Such a positive feed-forward cascade is consistent with the extensive enlargement of the perivascular immune niche after immunotherapy within responder animals, supporting the importance of this microenvironment subtype in the generation of highly localized and productive anti-tumor immune responses.

One additional point of consideration is the spatial patterning observed in the myeloid cell compartment with preferential association of DCs and activated Mfs near intra-tumoral blood vessels and deeper infiltration of the tumor nest by non-activated Mf populations. Such partitioning may, at least in part, be driven by chemotactic or adhesion properties of different myeloid cells and local guidance cues generated by other cell types (Ozga et al., 2021). In addition, direct access to glucose and other nutrients from the blood stream, coupled with reduced exposure to lactic acid within the tumor nest regions, is likely to spatially restrict innate cell function to the vicinity of blood vessels (Pearce and Everts, 2015; Wculek et al., 2019). Recent observations indicate increased consumption of glucose by DCs within tumors in comparison to other cell populations (Reinfeld et al., 2021), indicating increased metabolic dependence on perivascular localization for functional innate immune cells. Conversely, exposure to increased hypoxia within tumors can promote the generation of immunosuppressive Mfs, consistent with our and others’ findings on increased distances of non-activated Mfs from intra-tumoral vasculature (Murdoch et al., 2004; Doedens et al., 2010; Corzo et al., 2010; Broz et al., 2014; Henze and Mazzone, 2016; Huang et al., 2018). Thus, similar to adaptive lymphocytes, innate immune cell localization within the tumor may be a direct reflection of their functional properties, driven by local gradients of nutrients from local vasculature vs. exposure to suppressive factors generated by highly proliferative malignant cells. In this regard, therapies promoting vascular normalization have demonstrated efficacy when combined with immune targeted treatments (Huang et al., 2013; Schmittnaegel et al., 2017; Tian et al., 2017; Johansson-Percival et al., 2018; Huang et al., 2018; Munn and Jain, 2019; Lee et al., 2020; Zhang et al., 2021). Therefore, functional tumor vasculature appears to serve as a major organizational hub for spatially coordinated activities of innate and adaptive immune cells, both through improving local cellular recruitment and allowing normal metabolic and cellular functions.

Of note, responses of both MC38-CEA and KPC-CEA tumors to aPD-L1 monotherapy were limited in our studies, which likely reflects the relatively poor initial infiltration by CD8 T cells compared to ‘immunologically hot’ tumors (Feng et al., 2017; Mosely et al., 2017; Yu et al., 2018; Taylor et al., 2019). Both tumor types also displayed a general paucity of DC1s, which have been shown to be critical for promoting anti-tumor CD8 T cell responses in draining lymph nodes and within tumor tissues after checkpoint blockade therapy (Broz et al., 2014; Mikucki et al., 2015; Spranger et al., 2015, 2017). In this fashion, the balance of distinct DC subsets within tumors appears to set the tone of ongoing anti-tumor immune responses, which can then be potentiated by checkpoint inhibitors. The molecular factors dictating DC subset abundance in tumors are under investigation and appear to involve functional crosstalk of innate lymphocytes and tumor cells, as well as local generation of chemo-attractive factors (Böttcher and Reis e Sousa, 2018; Böttcher et al., 2018). While strategies to enhance DC1 infiltration into tumors are under exploration, bypassing such DC1 dependencies altogether may provide alternative strategies for immune control. Consistent with this, the use of bispecific antibodies which crosslink T cells with tumor antigens (i.e. TCB) likely promote the observed therapeutic effects independent of the DC1s’ cross-presentation abilities, although additional studies to evaluate the functional contributions of distinct innate populations in these settings are necessary.

In sum, our study provides enhanced resolution of the TME complexity and demonstrates existence of distinct immune microenvironments within tumors. We find evidence for the existence of the perivascular immune niche, suggesting that organization of immune cells within tumor tissues is dominantly shaped by the structural framework provided by local blood vessels. Finally, our findings demonstrate that implementation of quantitative imaging technologies has the potential to provide both insights into the mechanisms of immune cell function in tumors and generate companion and prognostic biomarkers associated with disease outcome, supporting continued development and use of such methods in cancer research.

## Author contributions

CRS, RS, LK, BEOP, EM, SC, MP and MYG edited manuscript.

## Lead contact and materials availability

Further information and requests for resources and reagents should be directed to and will be fulfilled by the lead contacts, Michael Gerner or Mario Perro. This□study did not generate new unique reagents.

## Materials and Methods

### Animals

Detailed methods can be found in Supplementary information.

### Cell lines

Detailed methods can be found in Supplementary information.

### Tumor studies

Six-to-nine week old huCEAtg mice were subcutaneously (s.c.) injected with 5×10^5^ MC38-huCEA (Robbins et al., 1991) or 3×10^5^ KPC-4662-huCEA cells (Lo et al., 2015) resuspended in RPMI medium with growth factor reduced Matrigel (Corning,354230) (1:1) in a total volume of 100ul in the right flank and tumor volume (1/2 [length X width^2^]) was measured 2-3 times per week using calipers. Mice were randomly assigned into different treatment groups based on tumor volume. Randomized mice (similar average tumor volume among all groups) were treated from day 19-21 post tumor cell injection twice per week intravenously (i.v.) with vehicle, or murine (muCEA-TCB) (surrogate version of RG7802, RO6958688) (2.5 mg/kg) and/or anti PD-L1, generated in house (RO7013159) (i.v. (first injection, 10mg/kg) or intraperitoneally (i.p.) for all further injections, 5mg/kg). Animals were sacrificed at day 27-29 post tumor cell injection.

Six-to-ten week-old BALB/c mice were s.c. injected with 5×10^5^ CT26.WT cell line resuspended in RPMI medium with growth factor reduced Matrigel (Corning,354230) (1:1) in a total volume of 100ul in the right flank and tumor volume measured twice per week using calipers. Mice were harvested 9 days after tumor injection for fixation and imaging.

Seven-week-old male B6 mice were s.c. injected with 5×10^5^ B16.F10.OVA.mCherry cell line resuspended in 1X Phosphate Buffered Saline (PBS,100ul) with growth factor reduced Matrigel (Corning,354230) (50ul) in a total volume of 150ul in the right flank and tumor volume measured twice per week using calipers. Mice were harvested 14 days after tumor injection for fixation and imaging.

### Tissue preparation and imaging

Harvested tumors were bisected and fixed with Cytofix (BD Biosciences) buffer diluted 1:3 with PBS for 12h at 4° C and then dehydrated with 30% sucrose in PBS for 12-24h at 4° C. Tissues were next embedded in O.C.T. compound (Tissue-Tek) and stored at −80° C. Tumors (MC38-hCEA, CT26, B16.F10.Ova mcherry) were sectioned on a Thermo Scientific Microm HM550 cryostat into 20μm sections. For the KPC-4662 huCEA model, tumors were fixed with 1:4 Cytofix solution for 21h at 4° C, embedded in 4% low gelling temperature agarose (Sigma) and sectioned into 70µm sections with a Vibratome (Leica 1200s).

Sample sections were prepared for imaging as previously described (Gerner et al., 2012). Briefly, sections were stained with panels of fluorescently conjugated antibodies (Table S1), cover-slipped with Fluoromount G mounting media (SouthernBiotech), and imaged on a Leica SP8 microscope using 40X 1.3NA (HC PL APO 40x/1.3 Oil CS2, for 20μm and 70μm sections) oil objective with type F immersion liquid (Leica, refractive index n_e_ = 1.5180). After acquisition, stitched images were compensated for spectral overlap between channels using the Leica Channel Dye Separation module in the Leica LASX software. For single stained controls, UltraComp beads (Affymetrix) were incubated with fluorescently conjugated antibodies, mounted on slides, and imaged with the same microscope settings as used to collect sample data. In all figures, for visual clarity, thresholds were applied to the displayed channel intensities.

### Image analysis and histo-cytometry

Image analysis and histo-cytometry was performed as described previously, with minor modifications (Gerner et al., 2012; Li et al., 2017, 2019; Stoltzfus et al., 2020b). Briefly, Imaris was used for initial image processing. Channel arithmetics were performed using either the default Imaris function or a customized ImarisXT extension, *Calebs_Multi_EQ_ChannelArithmetics_V3* (Table 1). Imaris was next used to segment individual cell objects or to generate spots representing different cells or tissue landmarks. After surface creation, the MFI for each imaged channel, as well as the volume, sphericity, and position of the cell objects were exported and concatenated into a single .csv file using the *Imaris_To_FlowJo_CSV_Converter_V6* MATLAB function, available online (Table 1). The combined .csv file was next imported into FlowJo (TreeStar) and the cell objects were classified into the indicated cell subsets according to the gating strategies shown in the respective figures.

### CytoMAP spatial analysis

Analysis of regions and spatial statistics was performed using CytoMAP, as described previously (Stoltzfus et al., 2020b). Details on CytoMAP analysis can be found in Supplementary information.

### Statistical analysis

No statistical method was used to predetermine sample size. The statistical significance of Pearson’s correlation was calculated using a Student’s t distribution for a transformation of the correlation.

## Data and code availability

All data are available upon request. Imaris extensions and other scripts used for histo-cytometry analysis are available for download at: https://gitlab.com/gernerlab/imarisxt_histocytometry CytoMAP software is available for download at: https://gitlab.com/gernerlab/cytomap

