## Supplemental Material for "Multi-Parameter Quantitative Imaging of Tumor Microenvironments Reveals Perivascular Immune Niches Associated with Anti-Tumor Immunity"

### *Supplementary Material*

#### Supplementary Materials and methods

##### Animals

Immunocompetent female human CEA transgenic (huCEA Tg) C57BL/6JTgN(CEAGe) 18FJP mice obtained under license agreement from Beckmann Research Institute of City of Hope (Clarke et al., 1998) at 6-9 weeks of age were maintained under specific pathogen free conditions with daily cycles of 12 hours light/ 12 hours darkness according to International (Federation of European Laboratory Animal Science Associations (Felasa)) and national (Gesellschaft fur Versuchstierkunde/Society of Laboratory Animal Science (GV-Solas) Tierschutzgesetz (TierSchG) guidelines. Experimental study protocol was reviewed and approved by local government authorities ((ZH227/17 and ZH193/2014). Animals were monitored on a regular basis and allowed to recover for 1-week post arrival to get accustomed to the new environment.

For CT26 tumor data presented in Supplementary Figure 4D, 6-10-week-old female Balb/c mice obtained from Charles River (Sulzfeld, Germany) were housed in specific pathogen-free conditions. The animal facility was accredited by the Association for Assessment and Accreditation of Laboratory Animal Care and all animal studies were performed in accordance to guidelines listed in Federation for Laboratory Animal Science Association and the German Animal Welfare Law. Experimental study protocol was reviewed and approved by Government of Upper Bavaria (Regierung von Oberbayern; license number: ROB-55.2-2532.Vet_03-15-41).

For B16.F10 data presented in Supplementary Figure 4D, 7-week-old male C57BL/6J mice obtained from The Jackson Laboratory were housed in specific pathogen-free conditions at Association for Assessment and Accreditation of Laboratory Animal Care- accredited animal facility at the University of Washington, South Lake Union campus. Experimental studies were performed in accordance to guidelines stated in the University of Washington Animal Care and Use Committee.

##### Cell lines and transduction

MC38-huCEA, mouse colon adenocarcinoma cell line engineered to express human CEA (Robbins et al., 1991) were maintained in RPMI medium supplemented with 10% FCS, 500ug/ml Geneticin (G418, Gibco). KPC-4662, murine pancreatic ductal adenocarcinoma cell line obtained from donating investigator (Dr. Vonderheide, University of Pennsylvania) (Lo et al., 2015) were transduced in house to express human CEA and maintained in DMEM with 1% L-Glutamine (Fisher Scientific) and 10% FBS (Corning). B16.F10 mouse melanoma cell line expressing chicken ovalbumin (B16.F10.Ova) and mcherry construct obtained from donating investigator (Dr.Oberst, University of Washington) were maintained in DMEM with 1% L-Glutamine (Fisher Scientific), 10% FBS(Corning), 1% Penicillin Streptomycin(VWR) and 1% Sodium Pyruvate (GE Healthcare).

##### CytoMAP spatial analysis

Analysis of regions and spatial statistics was performed using CytoMAP, as described previously (Stoltzfus et al., 2020b). Below is a brief discussion of the specific workflows used for the datasets described in this manuscript. The annotated cell surfaces for each dataset were loaded into CytoMAP by importing the corresponding paired .wsp and .fcs files after saving them in FlowJo. Once imported the following functions were used in CytoMAP to analyze the data.

*Cluster CEA and PD-L1 Spots*
CEA spot objects and PD-L1 spot objects were clustered using a self-organizing map. The mean fluorescent intensity of all 13 channels, standardized per sample, were used as the clustering parameters. Clusters were broadly grouped into subcategories based on their prevalence, channel intensity, and location in the image.

*Make surface*
To find the distance to the tumor border a surface was created around regions found by clustering neighborhoods using only the CEA and PD-L1 spot sub-types. This surface was created in CytoMAP using the *Make Surface* function. This function uses MATLAB's *alphaShape* function, with user defined parameters, to wrap the currently plotted points in a surface. This surface defines the $CEA^+$ Tumor region border shown in Figure 2B.

*Calculate distance*
This function was used to find the physical distance to the tumor border as well as various cell types shown in Figures 2C and S5.

*Raster scan neighborhoods*
This function was used to define neighborhoods for subsequent analysis shown in Figures 2, 3, 4, and 6. For the analysis presented here neighborhood analysis calculated the number of cells within a cylindrical volume in the tissue with a radius of 50 μm. The positions of the neighborhoods are evenly distributed throughout the tissue in a grid pattern with a distance between neighborhood centers of 25 μm.

*Classify neighborhoods into regions*
This function was used to define tissue regions. For all figures in this manuscript, we used the Global Composition (number of cells divided by the maximum number of cells in all neighborhoods across all samples). In this manuscript, the physical position of the neighborhoods was not used for region definition and the minimum of the Davies-Bouldin function was used to automatically determine the number of regions. If the global minimum of the Davies-Bouldin function was 2, then the next lowest minimum was chosen. The neighborhoods were clustered using the SOM function. For visualization purposes the names and colors of the regions were changed with the *Annotate Regions* function in CytoMAP. Some regions were combined for clarity. The region heatmaps and prevalence of the color-coded neighborhoods was plotted using the *Region Statistics* function in CytoMAP. The region heatmap plots the fold change in the number of objects per neighborhood within the indicated regions compared to the number of objects from all neighborhoods. The spatial distribution of the regions was visualized by generating a new figure in CytoMAP, plotting the positions of the neighborhoods, and selecting the regions for the 'c' axis to color-code the neighborhoods by region type.

*Reduce dimensions*
UMAP and t-SNE dimensionality reduction algorithms were used within CytoMAP to visualize tissue structure and complexity, as well as for treatment group comparisons. The Global

composition of the neighborhoods was used as the dimensions to be reduced. We used the MATLAB implementation of UMAP, provided by the Herzenberg Lab at Stanford University available for download at: https://www.mathworks.com/matlabcentral/fileexchange/71902-uniform-manifold-approximation-and-projection-umap.

#### Supplementary Tables

Table 1 Key Resources and Software

| REAGENT or RESOURCE | SOURCE | IDENTIFIER |
|---|---|---|
| Antibodies | | |
| All antibodies are listed in Table S5 | N/A | N/A |
| Chemicals, Peptides, and Recombinant Proteins | | |
| Tissue-Tek O.C.T. Compound | Electron Microscopy Sciences | Cat# 62550-01 |
| BD Cytofix fixation buffer | BD Biosciences | Cat# 554655 |
| PBS (pH 7.4) | Caisson Labs | Cat# PBL06-6X500ML |
| Triton-X-100 | Sigma-Aldrich | Cat# T-9284 |
| Bovine Serum Albumin | Sigma-Aldrich | Cat# A9576-50ML |
| Normal Mouse Serum | Jackson Laboratories | Cat# 015-000-120 |
| Tris buffer (1 M dilute to 0.1M) | Fisher Scientific | BP1756500 |
| Mix-n-Stain CF Dye Antibody Labeling Kits | Biotium | Cat# 92433-92339 |
| Agarose | Fisher Scientific | Cat# 16500500 |
| Immersion Oil, type F | Fisher Scientific | Cat# NC0586121 |
| EndoFit Ovalbumin 100mg | Invivogen | Cat# vac-nova-100 |
| Alhydrogel adjuvant 2% , 250mL | Invivogen | Cat# vac-alu-250 |
| Sucrose, ultrapure DNAse- and RNAse-free | VWR | Cat# 97061-432 |
| Software and Algorithms | | |
| CytoMAP | This Manuscript | https://gitlab.com/gernerlab/cytomap |
| Imaris extensions | This Manuscript | https://gitlab.com/gernerlab/imarisxt_histocytometry |
| Imaris | Bitplane | https://imaris.oxinst.com/ |
| LASX | Leica Microsystems | https://www.leica-microsystems.com/products/microscope-software/p/leica-las-x-ls/ |
| FlowJo | FlowJo, LLC | https://www.flowjo.com/ |
| Prism | GraphPad Software | https://www.graphpad.com/scientific-software/prism/ |

| | | |
|---|---|---|
| MATLAB | The MathWorks, Inc. | https://www.mathworks.com/products/matlab.html?s_tid=hp_products_matlab |
| Other | | |
| UltraComp eBeads Compensation Beads | Fisher Scientific | Cat # 01-2222-42 |
| PAP pen | Vector Laboratories | Cat# H-4000 |

#### Table S1. Antibody staining panels

| Ch | Antibody | Fluorophore |
|---|---|---|
| **MC38-CEA – Figures 1-3** | | |
| 1 | CD8 | BV421 |
| 2 | CD11c | BV480 |
| 3 | Foxp3 | eF506 |
| 4 | CD3 | Dyomics396xl |
| 5 | CD103 | CF633 |
| 6 | PD1 | CF514 |
| 7 | SIRPα | Atto 490LS |
| 8 | TCF1 | AF488 |
| 9 | CD206 | CF660 |
| 10 | MHCII | AF700 |
| 11 | PhosphoS6 | AF750 |
| 12 | CEA | CF555 |
| 13 | PDL1 | CF594 |
| **MC38-CEA – Figure 4** | | |
| 1 | CD8 | BV421 |
| 2 | CD11c | BV480 |
| 3 | CD3 | BV510 |
| 4 | MHCII | Dyomics396xl |
| 5 | CD103 | CF633 |
| 6 | TCF1 | AF488 |
| 7 | PD1 | CF514 |
| 8 | SIRPα | Atto 490LS |
| 9 | CEA | CF555 |
| 10 | PDL1 | CF594 |
| 11 | CD206 | CF660c |
| 12 | Ki67 | AF700 |
| 13 | CD31 | APC-Fire-750 |
| **CT26 Figure S4** | | |
| | CD11c | AF647 |
| | MHC-II | AF700 |
| | CD31 | AF488 |
| **B16.F10 Figure S4** | | |
| | CD11c | BV480 |
| | MHC-II | Dyomics396xl |
| | CD31 | AF488 |
| | CD3 | CF633 |
| **KPC-CEA Figure 5-6** | | |
| 1 | CD8 | BV421 |
| 2 | CD11c | BV480 |
| 3 | CD4 | BV570 |
| 4 | PD1 | CF532 |
| 5 | SIRPa | AF647 |
| 6 | MHC-II | AF700 |
| 7 | TCF1 | AF488 |
| 8 | CD31 | AF594 |
| 9 | FoxP3 | CF633 |
| 10 | CEA | CF555 |

#### Table S2. Image analysis and histo-cytometry

| **Channel Arithmetic** | | | |
|---|---|---|---|
| Channel name/Sample | Description | Equation | Samples |
| **MC38-CEA – Figures 1-3** | | | |
| Foxp3_Smoothed | | gaussian filter = 0.34um | All |
| CD3_Smoothed | | gaussian filter = 0.34um | All |
| TCF1_Smoothed | | gaussian filter = 0.34um | All |
| Phosphos6_Corrected | | (ch11-ch10).*(ch11>ch10) | All |
| Myeloid Channel V2 | Cd11c + MHC-II + CD103 + Sirpa + CD206 | (2.5.*ch2.*(ch2>5) + 1.5.*ch10.*(ch10>5) + 2.*ch5.*(ch5>10) + 1.5.*ch7.*(ch7>5) + ch9.*(ch9>10))./3.5 | C1, C7, D7 |
| Myeloid Channel V1 | Cd11c + MHC-II + CD103 + Sirpa + CD206 + PhosphoS6 | (2.5.*ch2.*(ch2>5) + 1.5.*ch10.*(ch10>5) + 2.*ch5.*(ch5>10) + 1.5.*ch7.*(ch7>5) + ch9.*(ch9>10) + 1.7.*ch11.*(ch11>10))./3.5 | A6, A5, B8, B6, B1, C9, D4, D3, D10 |
| Lymphocyte Channel V1 | (CD8 + CD3_Smoothed + PD1 + Foxp3_Smoothed) MHC2<60 | (1.7.*ch1.*(ch1>5) + 3.*ch15.*(ch15>5) + 1.7.*ch6.*(ch6>10).*(ch11<40) + 3.4.*ch14.*(ch14>5)).*(ch10<60)./1.5 | A6, A5, B8, B1, C9, D4, D3, D10 |
| Lymphocyte Channel V2 | (CD8 + CD3_Smoothed + PD1) MHC2<60 | (1.7.*ch1.*(ch1>5) + 3.*ch15.*(ch15>5) + 1.7.*ch6.*(ch6>10).*(ch11<40) ).*(ch10<60)./1.5 | D10, B6, C1, C7, D7 |
| **MC38-CEA – Figure 4** | | | |
| CD31 Corrected | | (ch12<30).*ch13 | All |
| Myeloid Channel | | ch2+ch4+ch8+(ch11.*(ch5<60)) | All |

| **Surface creation parameters** | | | | | | | | | |
|---|---|---|---|---|---|---|---|---|---|
| Surface Spots Name | Source Channel | Smoothing: Surface Detail (μm) | Background Subtraction: Diameter of Largest sphere | Absolute Intensity Threshold | Split Touching Objects: Split seed diameter (μm) | Quality Threshold | Voxel Number Threshold | | Diameter (Spots, μm) |
| **MC38-CEA – Figures 1-3** | | | | | | | | | |
| Lymphocyte Surface | Lymphocyte Channel | 0.67 | 10 | >30 | 6 | >10 | >500 | | N.A. |
| Myeloid Surface | Myeloid Channel | 0.67 | 20 | >25 (D7:>35) | 12.5 | 2 (D7: >6) | >50 | | N.A. |
| CEA Spots | CEA Channel | | none | | | >6 | | | 5 |
| PD-L1 Spots | PDL1 Channel | | none | | | >10 | | | 3 |
| **MC38-CEA – Figure 4** | | | | | | | | | |
| CD8 | CD8 Channel | 0.712 | 10 | >8 | 6 | >3 | >200 | | |
| CEA Spots | CEA Channel | | none | | | >25 | | | 15 |
| CD31 Surface | CD31 Corrected | 0.712 | 100 | >2 | 5 | >2 | >200 | | |
| Myeloid Surface | Myeloid Channel | 0.712 | 25 | >25 | 12 | >8 | >200 | | |
| **KPC-CEA Figure 5-6** | | | | | | | | | |
| CD8 Surface | CD8 Channel | 1.52 | Default | Default | Default | Default | >250 | | N.A. |
| CD4 Surface | CD4 Channel | 1.52 | Default | Default | Default | Default | >250 | | N.A. |
| Blood Vessels | CD31 Channel | 1.00 | 10 | Default | 4 | Default | >100 | | |
| CEA Spots | CEA Channel | | none | | | Default | | | 15 |

#### Table S3. Antibodies, related to reagents

| **Original MC38 Data** | | |
|---|---|---|
| CD3 –Dyomics396xl (clone 17A2) [Conjugated in house] | BioLegend | Cat# 100202 |
| CD8 –BV421 (clone 53-6.7) | BioLegend | Cat# 50-112-4973 |
| CD11c -BV480 (clone HL3) | BD | Cat# 565627 |
| CD103 -CF633 (goat polyclonal) [Conjugated in house] | R&D | Cat# AF1990 |
| CD206 -CF660 (clone MR5D3) [Conjugated in house] | R&D | Cat #MCA2235 |
| CEA -CF555 [Conjugated in house] | Roche | Cat# NA |
| Foxp3 -eF506 (clone FJK-16s) | eBioscience | Cat# 50-112-4973 |
| MHCII -AF700 (clone M5/114.15.2) | BioLegend | Cat# 107622 |
| PD1 -CF514 (goat polyclonal) [Conjugated in house] | R&D | Cat# AF1021 |

| | | |
|---|---|---|
| PD-L1 -CF594 (clone MIH5) | eBioscience | Cat# 14-5982-82 |
| PhosphoS6 rabbit (clone 2F9) | Cell Signaling | Cat# 4854S |
| SIRPα-biotin (clone P84) | eBioscience | Cat# 13-1721-82 |
| TCF1 –AF488 (clone C63D9) | Cell Signaling | Cat# 6444S |
| anti-rabbit-AF750 | Invitrogen | Cat# A-21039 |
| SA-ATTO490LS | ATTO-TEC | Cat# AD490LS-61 |
| **MC38 Vascular Panel Inclusions** | | |
| CD31 APC-Fire-750 (Clone Mec13.3) | BioLegend | Cat # 102514 |
| Ki-67 -AF700 (Clone 16A8) | Biolegend | Cat # 652420 |
| CD3 -BV510 (Clone 17A2) | Biolegend | Cat # 100234 |
| MHC -II Dyomics396xl (clone M5/114.15.2) [Conjugated in house] | Biolegend | Cat # 107602 |
| **KPC Data** | | |
| CD8 -BV421 (clone 53-6.7) | Biolegend | Cat # 100738 |
| CD11c -BV480 (clone N418) | BD Biosciences | Cat # 746392 |
| CD4 -BV570 (clone RM4-5) | Biolegend | Cat # 100542 |
| PD1 (polyclonal) | R&D | Cat # AF1021 |
| CF532 | Biotium | Cat # 92290 |
| SIRPa -AF647 (clone P84) | Biolegend | Cat # 144028 |
| MHC-II -AF700 (clone M5/114.15.2) | Biolegend | Cat # 107622 |
| TCF1 -AF488 (clone C63D9) | Cell Signaling Technologies | Cat # 6444S |
| CD31 -AF594 (clone MEC13.3) | Biolegend | Cat # 102520 |
| FoxP3 (clone D6O8R) | Cell Signaling Technologies | Cat # 12653 |
| CF633 | Biotium | Cat # 92257 |
| CEA (in-house T84.66) | in-house | N.A. |
| CF555 | Biotium | Cat # 92254 |

#### Supplementary figures

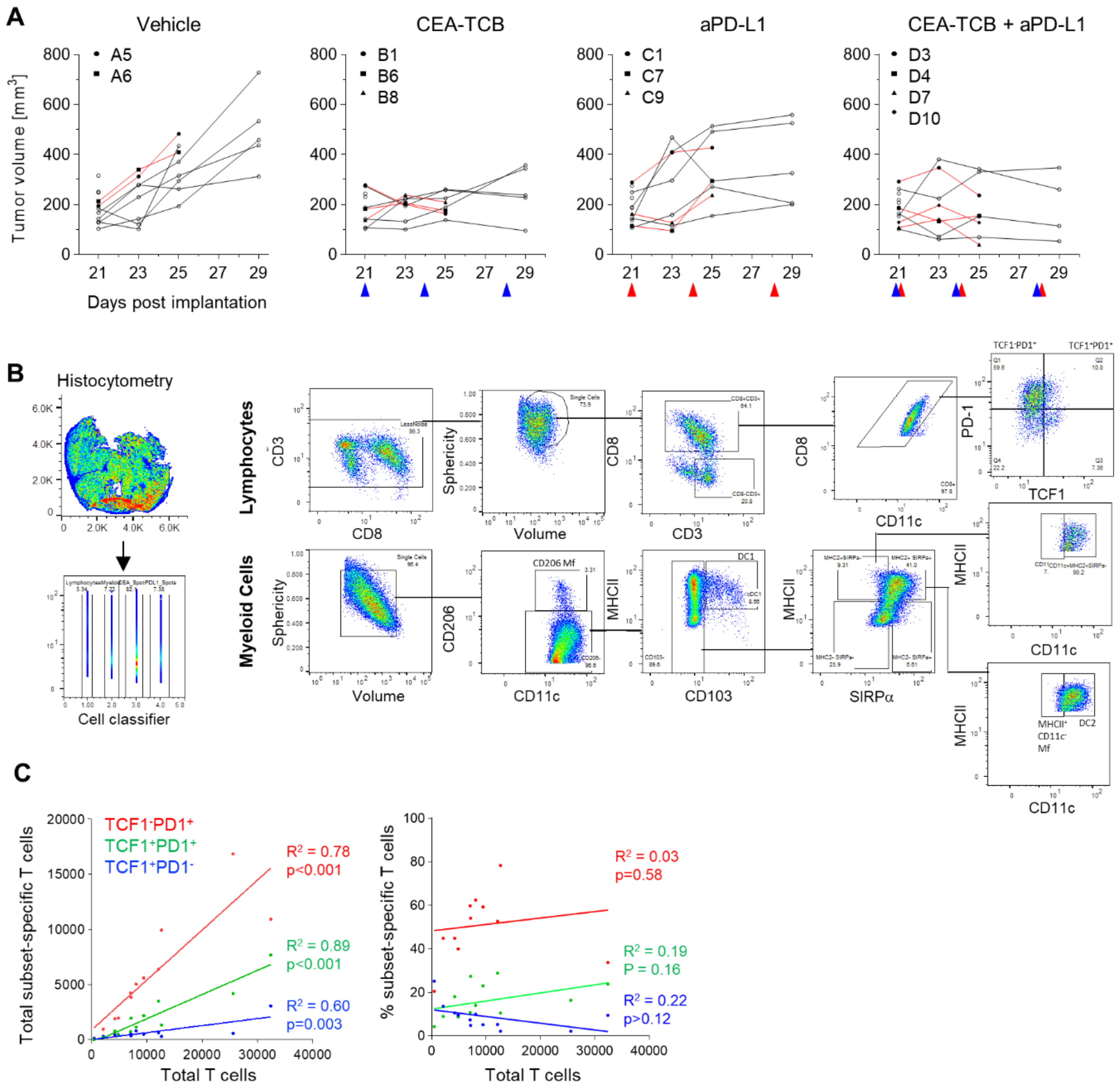

**Supplementary Figure 1: Histocytometry based phenotyping of immune cells from the MC38-CEA tumor microenvironment. (A)** MC38-CEA tumor volume is shown versus days since implantation. Arrows denote treatment timepoints. Tumors used for subsequent imaging are indicated by red lines and were sacrificed 4d post initiation of treatment. **(B)** The histocytometry gating strategy used to phenotype select immune cell populations in the MC38-CEA tumors. Separate surface objects were created for lymphocyte and myeloid cell types. **(C)** Correlation between the total number of CD8 T cells and either the total number (left) or percent (right) of the indicated T cell subset.

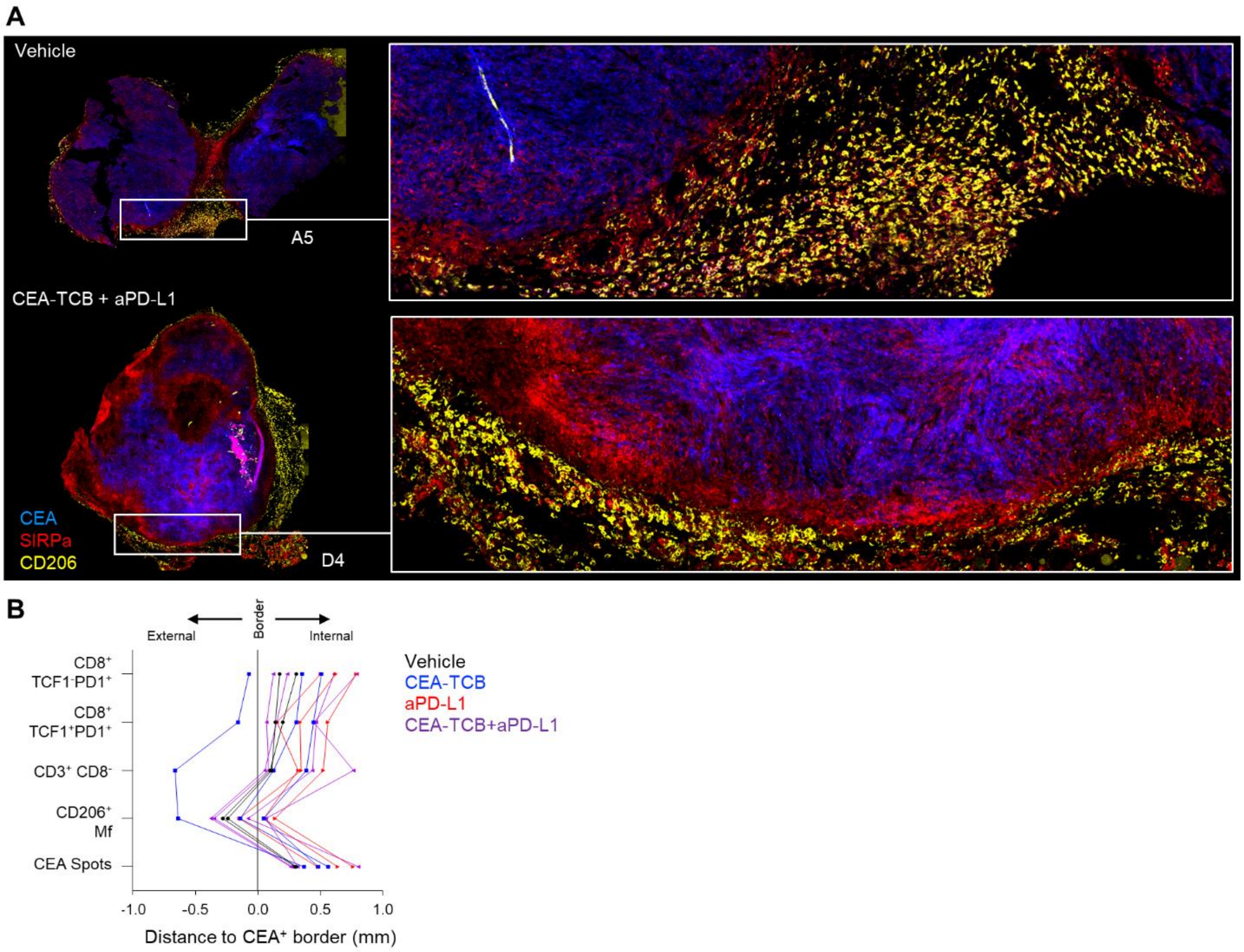

**Supplementary Figure 2: Defining MC38-CEA tumor border regions using CD206+ macrophages. (A)** Confocal images of selected tumors from the control and CEA-TCB plus aPD-L1 groups highlight the association between CD206+ macrophages and the external border region of the MC38-CEA tumor. **(B)** Average distance of the indicated cell populations to the CEA$^{+}$ tumor border plotted for all samples. Each connected line represents a single tumor cross-section.

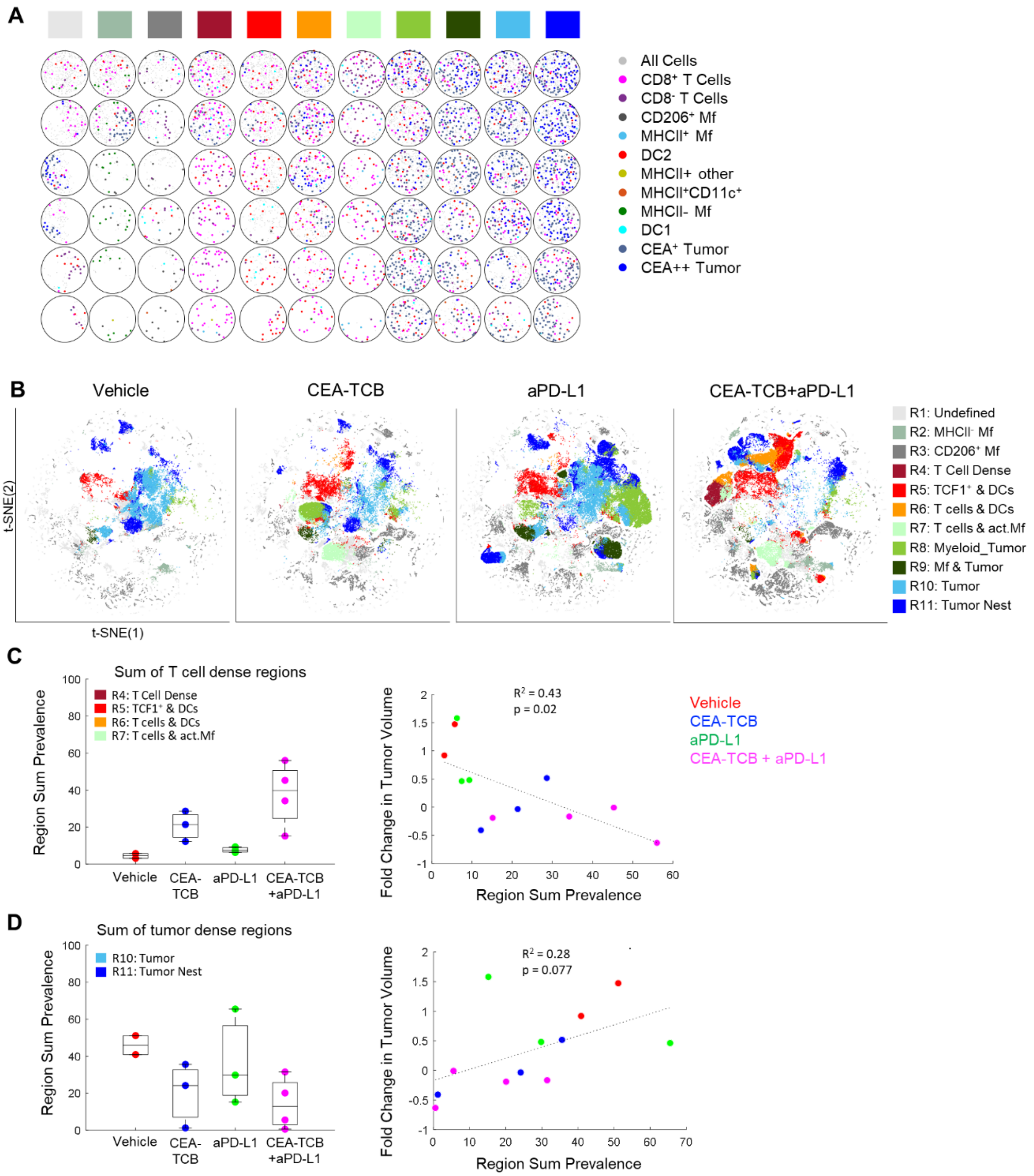

**Supplementary Figure 3: CytoMAP based analysis of MC38-CEA tumors. (A)** Positional plots of objects within select neighborhoods from each neighborhood type in MC38-CEA tumors. Color code for each neighborhood type (top) is defined in Fig. 3A. **(B)** t-SNE plots of the color-coded neighborhoods from all samples in each treatment group displaying the heterogeneity of tissue regions. **(C)** Prevalence of combined T cell dense regions (left) and their linear regression with tumor volume across all samples by group (right). **(D)** Prevalence of combined tumor dense regions (left) and their linear regression with tumor volume across all samples by group. All the above plots were generated as described in Fig S1A; n=2 for control, n=3 for CEA-TCB and aPDL1 and n=4 for CEA-TCB+aPD-L1 group.

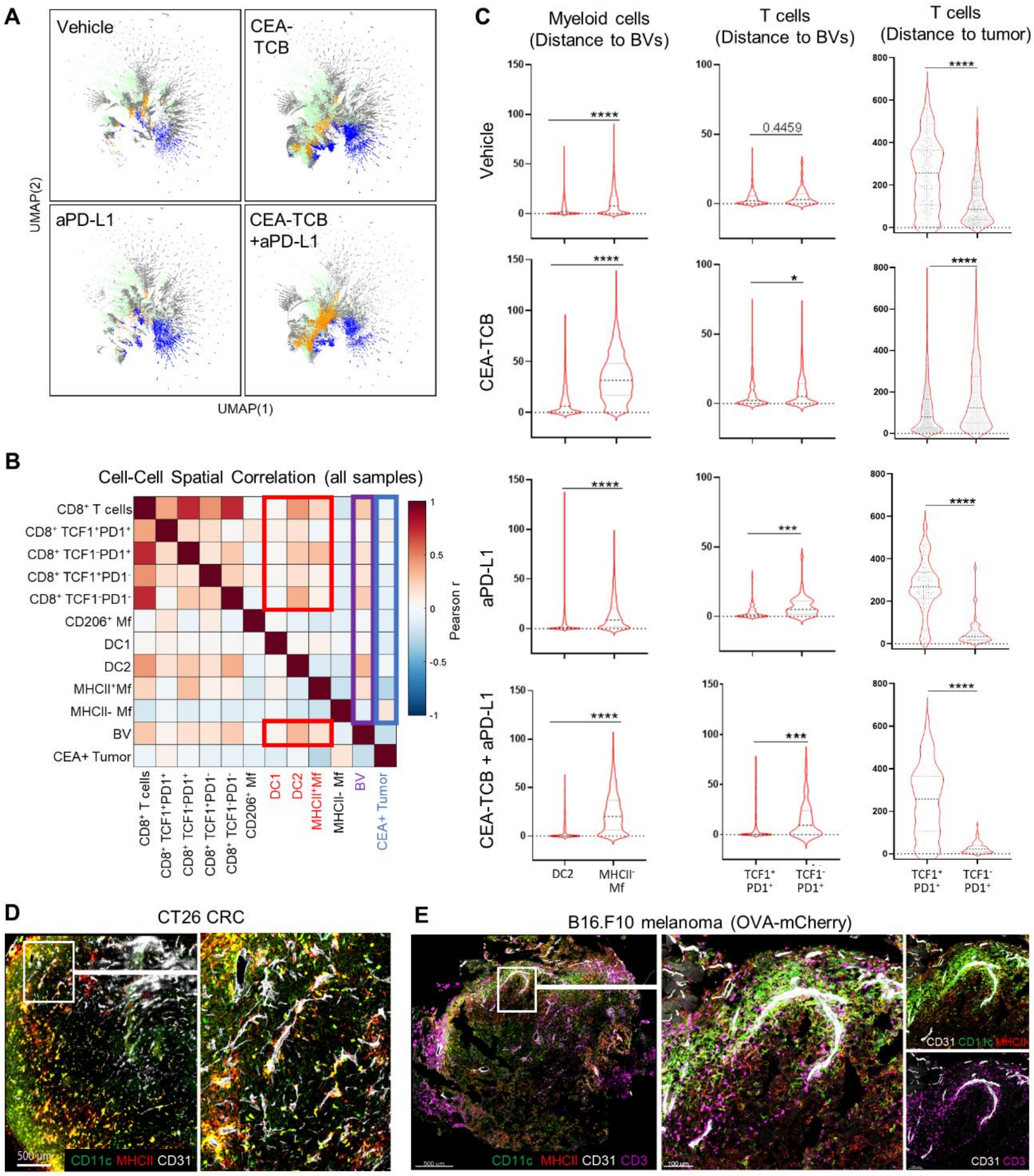

**Supplementary Figure 4: Analysis and prevalence of tumor associated perivascular immune niches. (A)** UMAP plots of the neighborhoods color-coded by region type as defined in Fig. 4B. **(B)** Heatmap of Pearson correlation coefficients between number of cells per neighborhood for each cell population pair within 50 µm neighborhoods from all imaged samples. **(C)** Violin plots showing the distance of myeloid cells or T cells to the nearest BV or CEA$^{+}$ tumor signal. All plots were generated from samples collected at d25, n=1 per group. **(D)** Multiplex confocal images visualizing the associations of DCs (CD11c$^{+}$MHCII$^{+}$) with CD31$^{+}$ BVs in CT26 CRC tumor samples. **(E)** Multiplex confocal images visualizing the associations of DCs (CD11c$^{+}$MHCII$^{+}$) and CD3 T cells with CD31$^{+}$ BVs in B16.F10 tumor samples.

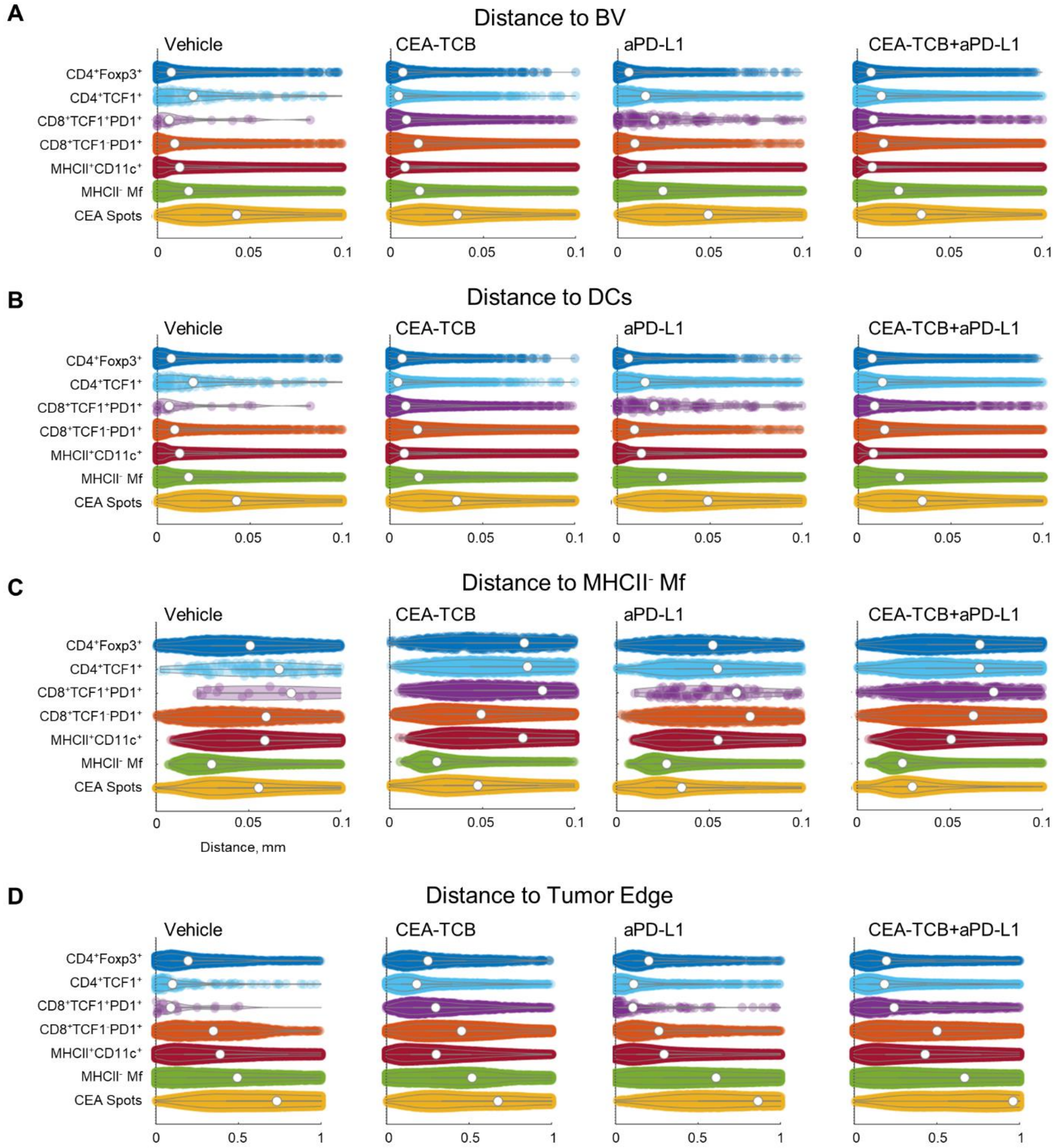

**Supplementary Figure 5: CytoMAP based distance analysis in KPC-CEA tumors**. Violin plots displaying the distance of the indicated cell types to the nearest **(A)** BV object, **(B)** DC, and **(C)** MHCII⁻ non-activated Mf. **(D)** tumor edge, as defined manually in KPC-CEA tumors. Data in A-D plot all cells from all samples for each treatment group. Data represent one experiment with 3 samples per group.

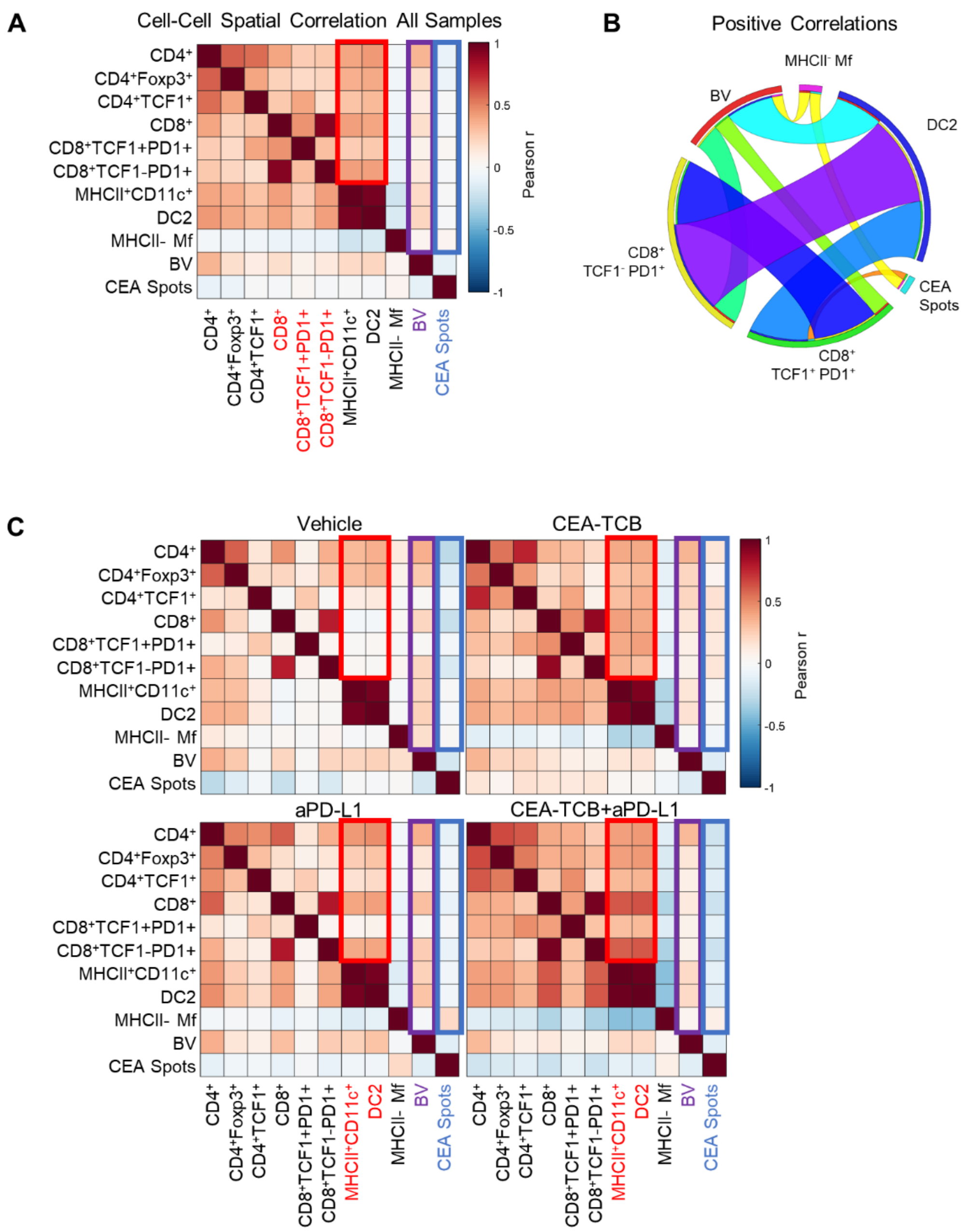

**Supplementary Figure 6: Spatial correlation patterns in the KPC-CEA tumor microenvironment. (A)** Heatmap displaying the spatial correlation between the number of cells per neighborhood for each cell population pair from all imaged samples in the KPC-CEA tumor. **(B)** Circos plots of the scaled and rounded positive Pearson correlation between the number of cells per neighborhood across all samples for the indicated cell types. Plots were generated using www.circos.ca. The colors in the circos plot are auto-generated. **(C)** Heatmaps displaying the spatial correlation between the cell density for each cell population pair for all samples by treatment group. Data from the KPC-CEA tumor models are described in Figure 5.

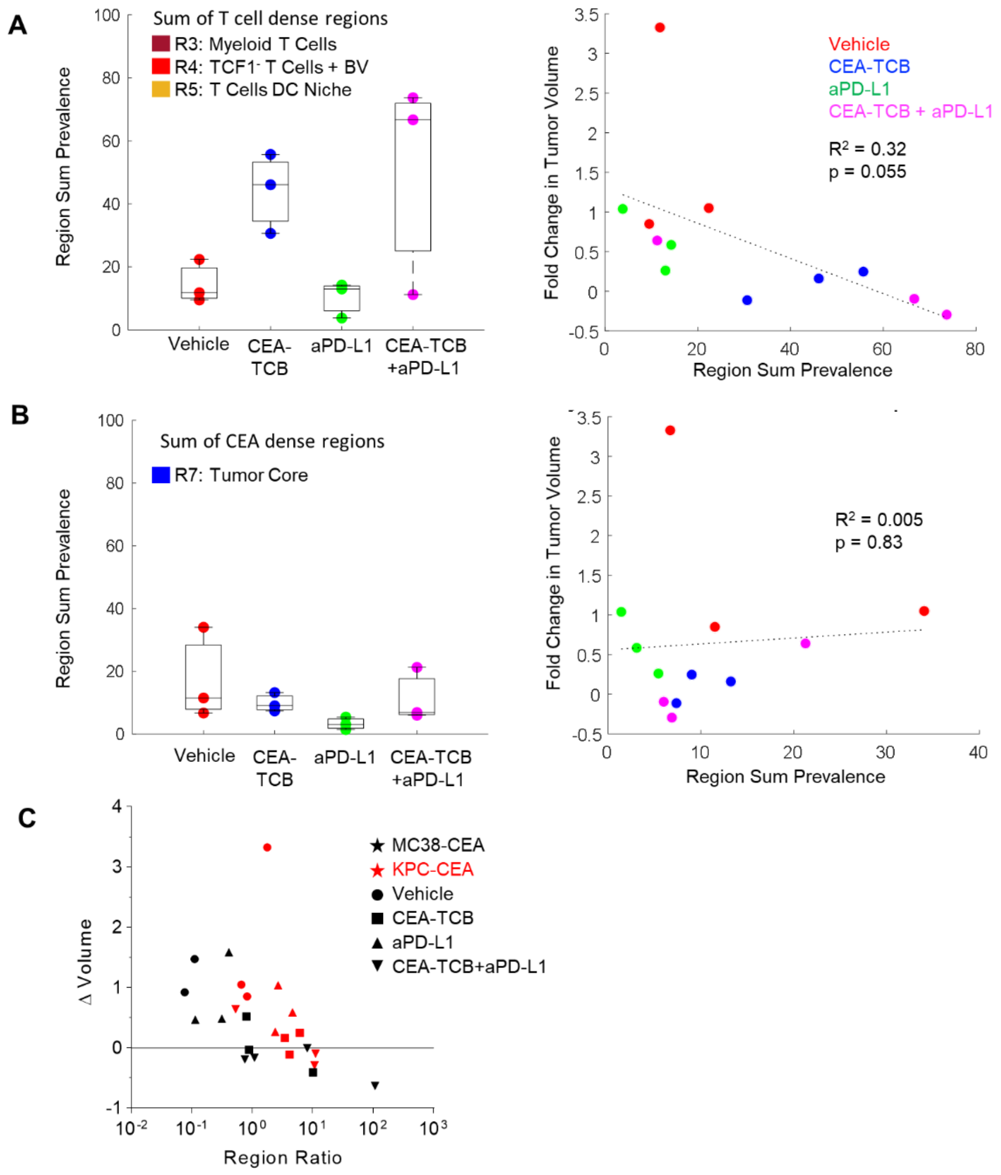

**Supplementary Figure 7: Association between KPC-CEA tumor regions and tumor regression. (A)** Prevalence of combined T cell dense regions (left) and their linear regression with fold change in tumor volume across all samples by treatment group in KPC-CEA tumors. **(B)** Prevalence of combined tumor dense regions (left) and their linear regression with tumor volume across all samples by treatment group in KPC-CEA tumors. **(C)** Combined plot of the ratio data also shown in Figures 3G and 6F.
